# Atomistic Basis of Force Generation, Translocation, and Coordination in a Viral Genome Packaging Motor

**DOI:** 10.1101/2020.07.27.223032

**Authors:** Joshua Pajak, Erik Dill, Emilio Reyes-Aldrete, Mark A. White, Brian A. Kelch, Paul Jardine, Gaurav Arya, Marc C. Morais

## Abstract

Double-stranded DNA viruses package their genomes into pre-assembled capsids using virally-encoded ASCE ATPase ring motors. We present the first atomic-resolution crystal structure of a multimeric ring form of a viral dsDNA packaging motor and characterize its atomic-level dynamics *via* long timescale molecular dynamics simulations. Based on the results, we deduce an overall packaging mechanism that is driven by helical-to-planar transitions of the ring motor. These transitions are coordinated by inter-subunit interactions that regulate catalytic and force-generating events. Stepwise ATP binding to individual subunits increase their affinity for the helical DNA phosphate backbone, resulting in distortion away from the planar ring towards a helical configuration, inducing mechanical strain. Subsequent sequential hydrolysis events alleviate the accumulated mechanical strain, allowing a stepwise return of the motor to the planar conformation, translocating DNA in the process. This type of helical-to-planar mechanism could serve as a general framework for ring ATPases that exhibit burst-dwell dynamics.

## Introduction

The Additional Strand, Conserved Glutamate (ASCE) superfamily is an ancient and ubiquitous class of NTPases, encompassing subfamilies such as AAA+ motors, RecA-/FtsK-like ATPases, and ABC transporters^1^. These motors convert energy from NTP binding and/or hydrolysis into mechanical work, and typically perform biological segregation tasks such as proton transport, chromosomal segregation, DNA or RNA strand separation, and protein degradation. Double-stranded DNA (dsDNA) viruses, such as herpes-, adeno-, and pox viruses, as well all tailed bacteriophages, encode for ASCE segregation motors that they use to package their genomes into preformed procapsids during virus replication^2-4^. Among ASCE ATPases, viral packaging motors generate particularly high forces (>50 pN) to overcome the entropy loss, electrostatic repulsion, and DNA stiffness that oppose DNA confinement^5-8^. Thus, viral dsDNA packaging motors provide a unique window into the mechanochemistry of force-generation found in this broad class of molecular motors.

The relatively small size and simplicity of φ29-like phages has facilitated advanced genetic, biochemical, and structural studies^3^. For example, all components of the φ29 packaging system have been thoroughly characterized, and a robust highly efficient *in vitro* DNA packaging system has been developed^9^. Furthermore, atomic resolution structures of all individual φ29 motor components are available^10-13^ and medium resolution structures of motors assembled on capsids in various stages of assembly and/or packaging have been determined^13-18^. These results indicate that the DNA packaging motor consists of a dodecameric portal protein, a pentameric prohead RNA (pRNA), and a pentameric ATPase (gene product 16; gp16) that assemble as co-axial rings at a unique vertex of the φ29 capsid.

Due in part to this experimental accessibility, single-molecule force spectroscopy (SMFS) experiments have provided valuable information on force-generation and dynamics of viral packaging motors. High-resolution measurements showed that the motor packages DNA in 10 bp “bursts” comprised of four 2.5 bp sub-steps, each coupled to ATP hydrolysis and phosphate release. DNA translocation bursts are followed by a relatively long “dwell” wherein DNA translocation pauses while each ADP is sequentially exchanged for ATP to reset the motor for the next burst^19-21^.

Whereas genetic, biochemical, structural, and single-molecule studies have provided significant insights into the mechanochemistry of the φ29 packaging motor, the molecular basis of force-generation and coordination remains unresolved for any viral DNA packaging motor. Given the multi-component nature of the motor, it is difficult to determine how such coordination arises in the absence of high-resolution quaternary structural information. Likewise, it is difficult to determine the mechanisms of force generation in the absence of atomistic dynamic information. Hence, we determined the first high-resolution structure of a functional assembly of a φ29-like (asccφ28) ATPase and probed its dynamics *via* long-timescale molecular dynamics (MD) simulations. Together, these data resolve fundamental questions regarding inter-subunit coordination and force-generation, and enabled development of an atomistic model of viral DNA packaging wherein the motor transitions between helical and planar configurations to efficiently package DNA.

## Results

Despite extensive efforts, it has not been possible to assemble the functional ring-form of the packaging ATPase from bacteriophage φ29 detached from procapsids. This is not surprising, since the φ29 ATPase only assembles functional rings by virtue of binding to the procapsid^17^. In contrast, a close homolog of the φ29 ATPase from the related bacteriophage asccφ28 (gp11) has been shown to form highly soluble functional rings; extensive analytical ultracentrifugation and small-angle X-ray scattering experiments show that the ATPase forms pentameric rings in solution^22^. The 45% sequence similarity between the two proteins assures their structures will be nearly identical. Thus, we used asccφ28 gp11 for crystallographic structure determination.

### X-ray crystal structure determination

Cloning, protein purification, kinetic analysis, AUC, SAXS, negative stain TEM, and preliminary crystallization of gp11 has been previously reported^22^. Briefly, two crystal forms of gp11 were obtained: 1) tetragonal crystals belonging to space group P4_3_2_1_2 and 2) trigonal crystals belonging to space group P3_2_2_1_. Single-wavelength anomalous dispersion (SAD) was used to obtain initial experimental phases for the P3221 crystals grown from seleno-methionine labeled protein (**Table S1**). The final refined structure was used as a molecular replacement search model to phase the data from the P4_3_2_1_2 space group.

### Quaternary structure of gp11

Both the tetragonal and trigonal crystal forms had similar pentameric rings in their crystallographic asymmetric units despite having substantially different packing environments (**Fig. 1**), indicating that the observed pentamer is the biological assembly. This stoichiometry is consistent with previously reported biochemical and biophysical analysis that indicated gp11 forms pentameric rings in solution^22^. Of note, the kinetic parameters of ATP binding and hydrolysis by isolated gp11 rings are similar to the parameters obtained for φ29 and other bacteriophage DNA packaging motors, but only once these other ATPases are assembled as functional rings on their respective procapsids^22^. In the absence of procapsids, other packaging motors negligibly hydrolyze ATP, presumably since they are in monomeric form and therefore cannot efficiently bind or hydrolyze ATP (see also below). Additionally, the arrangement of the subunits is similar to the recent structure of the φ29 particles stalled during packaging^18^, with the notable difference that the φ29 ATPase structure is helical rather than planar. Hence, the pentameric stoichiometry of gp11 observed here reflects a functional assembly during DNA packaging.

**Figure 1:**
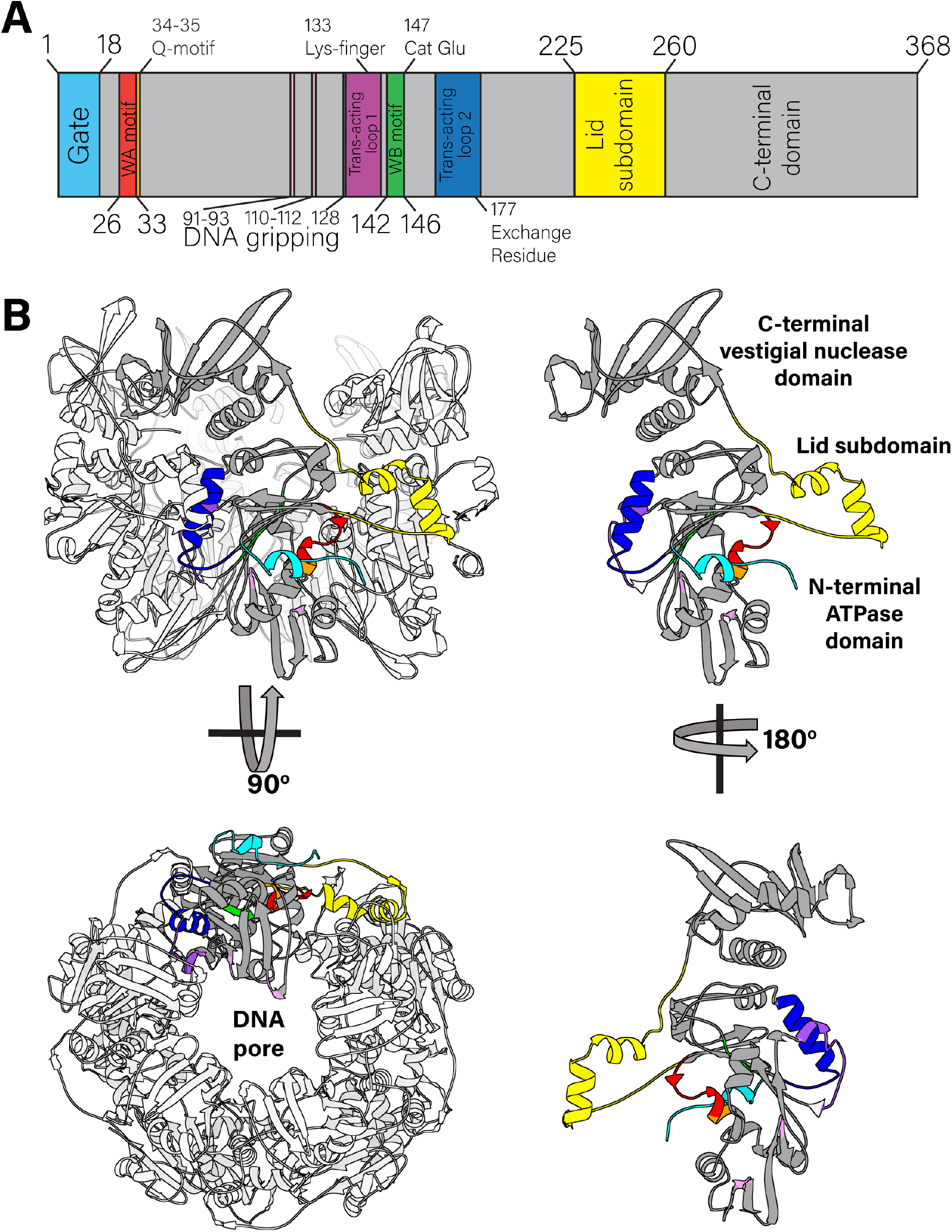
Quaternary and tertiary structure of the asccφ28 packaging ATPase. **A.** Sequence of the packaging ATPase contains canonical ASCE motifs, such as the Walker A and Walker B motifs. **B.** Structure of the pentamer (left), with a single subunit highlighted in gray and its lid subdomain highlighted in yellow is shown from side and end-on views (top and bottom panels, respectively). The lid subdomain mediates most inter-subunit contacts. The monomer in isolation (right) is color coded to match the motif classification in panel A, and is shown in two side views, from the exterior of the motor (top) and the interior of the lumen (bottom).

### Tertiary structure of gp11

Individual subunits within the ring are organized into two globular domains connected by a linker domain, each of which is nearly identical to the corresponding domains of the φ29 gp16 (**Fig. S1**)^18^. Like φ29 gp16, the N-terminal domain adopts the canonical ASCE ATPase fold, while the C-terminal domain is a “vestigial nuclease domain”^11^. The linker domain (residues 225 to 260) adopts a helix-loop-helix fold, again like φ29 gp16 and reminiscent of the lid subdomain identified in other ASCE ATPases such as AAA+ and helicases^1^.

### ATPase active site: *cis*-contributions

While the three structures described here were determined in the absence of nucleotide, the active sites of ASCE ATPases are well characterized and can be accurately identified. In the solved structures, the ASCE ATPase active site is situated between two subunits, such that both subunits participate in catalysis. The *cis-*acting side of the ATP-binding interface resides on one edge of the central beta-sheet of the Rossmann fold and includes canonical Walker A (26-GGRGVGKT-33) and Walker B (142-YLVFD-146) motifs.

Two distinct conformations of the Walker A motif are apparent in our crystal structures (**Fig. S2**). One conformation binds either a SO_4_^2-^ or PO_4_^3-^ ion in the canonical β-phosphate position. The ion forms hydrogen bonds with the backbone of the Walker A motif in lieu of a β-phosphate, typically provided by ATP or ADP, which explains why the Walker A backbone adopts a conformation similar to a typical nucleotide-bound ATPase configuration. However, because the structure was solved in absence of nucleotide or nucleotide analog, it lacks any interactions mediated by the α- and γ-phosphates, ribose sugar, or adenosine base. Thus, this structure likely represents a conformation with mixed characteristics from both the apo and nucleotide-bound configurations.

In contrast, the conformation observed in the iodine-bound structure shows that an iodine atom sits in a hydrophobic pocket that is occupied by Val30 in the other structures. As a result of displacing Val30, the Walker A motif adopts an extra helical turn in the phosphate-binding loop. This conformational change excludes the possibility of binding nucleotide since: 1) the sidechain of Val30 is in the center of the active site, occluding nucleotide, and 2) the Walker A backbone amide groups are repositioned such that they can no longer hydrogen bond with the β-phosphate. Initially, we assumed that the nucleotide-blocking valine was an artifact of heavy atom derivatization. However, subsequent structural analysis and MD simulations indicated that the two conformations of this loop may reflect an ability to switch between nucleotide accepting and occluding conformations, providing a mechanism for regulation of ATP binding and ADP release (see below).

### ATPase active site: *trans*-contributions

The *trans*-acting side of the subunit interface consists primarily of two helical segments (residues 129-139 and 161-178) that reside on the side of the central Rossmann fold opposite the *cis*-contributing elements. These helices position polar- and positively-charged residues in the ‘neighboring’ active site that contribute to ATP binding and hydrolysis, as well as to phosphate and ADP release (**Fig. 1**). Notably, Arg177 is positioned in the active site of all three crystal structures. Arg177 in asccφ28 corresponds to Arg146 in the φ29 ATPase, which had previously been identified as a *trans*-acting arginine finger^12,23^. Arginine fingers are ubiquitous in ring ATPases and are generally believed to catalyze ATP hydrolysis by stabilizing the transition state^24^. However, the sidechain of Arg177 is not optimally positioned to coordinate the expected position of the γ-phosphate of ATP (**Fig. S2**). Thus, for Arg177 to act as the arginine finger and catalyze hydrolysis *in trans,* the interface would require significant rearrangements.

A different positively charged residue, Lys133, is better positioned to interact *in trans* with the expected position of the γ-phosphate of ATP (**Fig. S2**). Such interaction would require significantly fewer structural rearrangements. Further, Lys133 corresponds to the arginine finger identified in the bacteriophage P74-26 packaging ATPase^25^, indicating that Lys133 may function as a “lysine finger.” We note that the use of a lysine to catalyze hydrolysis *in trans* instead of the “traditional” arginine finger has been observed in both RecA^26^ and DnaC^27^ ATPases, and thus there is precedence for a lysine finger in ATPase ring motors.

### Phosphate binding and regulation of hydrolysis

To further understand how subunits around the ring bind ATP, and how this binding might modulate DNA gripping, we performed long timescale MD simulations of the ATPase ring. For the starting structure, Mg^2+^-ATP was positioned into each subunit according to the structure of the BeF3-ADP-bound phage P74-26 packaging ATPase^25^. Additionally, a 30-bp, B-form dsDNA was placed in the central pore of the pentamer. The structure was equilibrated and its equilibrium dynamics were sampled *via* MD simulations for 2.4 μs. The simulations predicted that Mg^2+^-ATP binds to the *cis*-acting side of the inter-subunit active site *via* canonical interactions with the Walker A motif: 1) the β-phosphate of ATP forms several hydrogen bonds with the backbone nitrogens of the Walker A motif; 2) the critical P-loop Lys32 coordinates the β- and γ-phosphates; and 3) Thr33 chelates the Mg^2+^ ion (**Fig. 2A**). The *cis-*acting Walker B motif residues engage in similar conserved canonical interactions: 1) Asp146 chelates the Mg^2+^ ion through a water molecule, and 2) Asp146 hydrogen bonds to the Walker A Thr33, which has been previously predicted to be a key interaction that helps close the active site as part of the tight-binding transition^28,29^.

**Figure 2:**
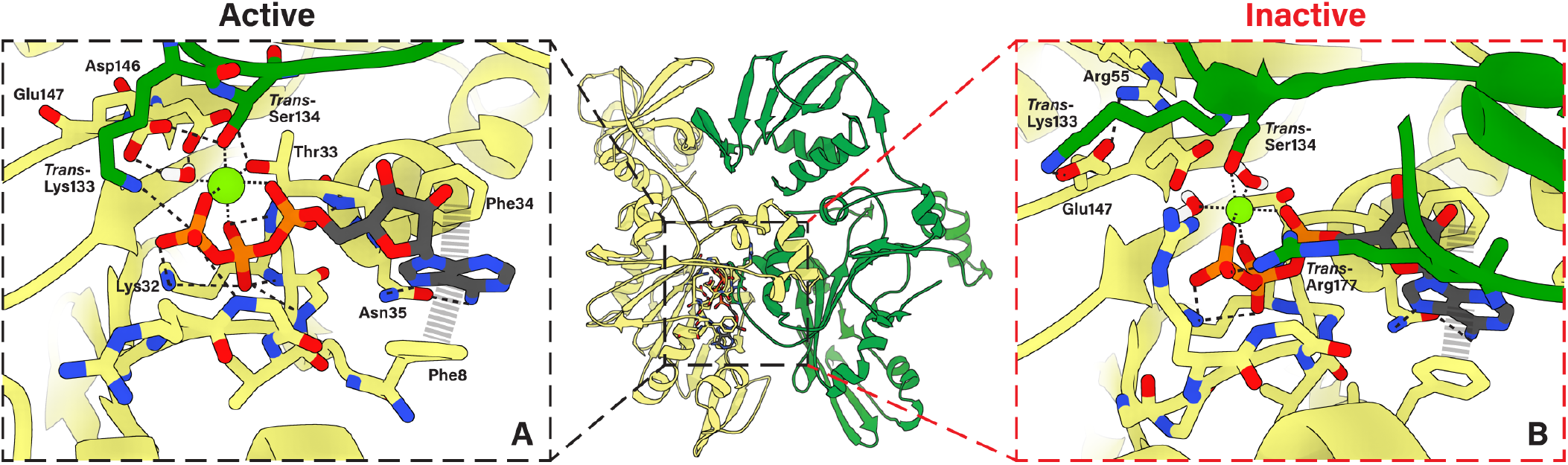
ATP-binding poses predicted from MD simulations. **A.** Active ATP-binding pocket contains canonically predicted interactions. The cis-acting (yellow) subunit’s Walker A motif backbone NH groups bind the β-phosphate, Lys32 binds the γ- and β-phosphates, and Thr33 chelates Mg^2+^. Downstream of the Walker A motif, Phe34 and Asn35 function as the Q-motif, and bind the adenosine with help from N-terminal gate Phe8. The Walker B motif Asp146 and catalytic Glu147 isolate a single water molecule, which chelates Mg^2+^ (green sphere). Residues donated in trans from the neighboring subunit (green) also participate in ATP-binding. Notably, Lys133 is seen interacting with the γ-phosphate and Ser134 chelates Mg^2+^. **B.** Inactive pose maintains many of the same interactions as the active pose, with a few key differences. Arg177 is now donated in trans to interact with the γ-phosphate, while Lys133 interacts with catalytic Glu147 away from the γ-phosphate. This interaction is stabilized by cis-acting Arg55.

While the *cis*-acting interactions were expected, the simulations further predicted that the *trans*-acting residues donated from the neighboring subunit form a tight hydrogen bonding network with the *cis* acting Walker motifs that is centered around the γ-phosphate (**Fig. 2A**). *Trans*-acting Lys133 interacts with the γ-phosphate of ATP, consistent with Lys133 functioning analogously to “arginine fingers” decribed in other systems. Such an interaction would help polarize the P-O bond, and stabilize the negative charge on the transition state^24^. Further, the residue immediately downstream of Lys133, Ser134, chelates the Mg^2+^ ion *in trans*. This lysine-serine pair is distinct from SRC motifs found in many AAA+ motors that contain an arginine finger^30^. In the AAA+ SRC motif, the serine residue is upstream of the *trans*-acting catalytic residue, and does not chelate Mg^2+^. Thus, to the best of our knowledge, our structure and simulations predict a new motif for *trans*-activated catalysis in ASCE enzymes.

The simulations also predicted that the *cis-*acting Walker B catalytic Glu147 hydrogen bonds to this *trans*-acting Ser134. The consequence of these interactions is a tight hydrogen bonding network centered around a single water molecule. The hydrogen atoms of this caged water hydrogen bond with the oxygens on the Walker B 146-DE-147 carboxylate groups, whereas the oxygen atom chelates the Mg^2+^ ion. Such a configuration would polarize electron density onto the oxygen atom. Hence, the water molecule is primed for deprotonation by Glu147, with the resulting nucleophilic hydroxide ion poised for attack at the γ-phosphate as described above. It is interesting to consider the conservation of the DE pair in light of our structures and simulation; the geometry of the active site ensures that the glutamate is positioned such that deprotonation leaves the lone pair of electrons on the nucleophilic hydroxide pointing directly at the phosphate target. If the positions of Asp and Glu were switched, the hydroxide ion would not be optimally oriented for nucleophilic attack. Indeed, DE switch mutations in viral packaging ATPases typically abrogate ATPase activity^15,31^.

In addition to the hydrolysis-competent active site described above, our simulations suggests that the binding interface can also adopt an “inactive” conformation. This conformation positions the catalytic Glu147 carboxylate group away from the γ-phosphate and towards *cis*-acting Arg55. We also found that while Ser134 still chelates Mg^2+^ *in trans* in the inactive conformation, Lys133 no longer interacts with the γ-phosphate and instead interacts with Glu147, helping Arg55 stabilize the inactive pose of the binding interface. The resuting interaction between Glu147 and Arg55 is analogous to the “glutamate switch” interaction found in AAA+ enzymes, and is indicative of a binding interface that catalytically incompetent^32^. In these other systems, the glutamate switch regulates the timing of hydrolysis, thus we suspect a similar role in viral DNA packaging motors. Indeed, a similar interaction is found in crystal structures of the φ29 and Sf6 packaging ATPases with bound nucleotide (**Fig. S3**).

### Adenosine binding and control over active site accessibility

The adenosine base of ATP binds similarly in both the “inactive” and “active” interfaces (**Fig. 2A,B**). An aromatic residue immediately downstream of the canonical Walker A motif, Phe34, π-stacks with the adenosine base. The adenosine forms bidentate hydrogen bonds with the next residue, Asn35. A pairing of an aromatic residue followed by a carboxamide has been previously identified as the “Q motif” in DEAD-box RNA helicases and viral packaging ATPases. The pair’s function was implicated in binding the adenosine ring in the λ phage packaging ATPase^33^.

On the other side of the adensoine base, the N-terminal loop repositions itself so that Phe8 π-stacks on the side of the adenosine opposite of Phe34. This contrasts with the position of the N-terminal loop in the crystal structure, which positions Phe8 farther away from the Walker A motif. Thus, the N-terminal loop forms a gate that can either open to allow nucleotide exchange, or close to tightly bind ATP. Similar π-stacking sandwiches flanking either side of adenosine have been observed in solved structures of other ATPases, such as *E. coli* MutS and the Human Catalytic Step I Spliceosome^34,35^. This suggests that similar active-site gating mechanisms may be present in other ring ATPases. In the closely related bacteriophage φ29, the N-terminal loop makes critical contact with pRNA^18^. Although gp11 was solved in absence of pRNA, the positions of all pRNA-contacting reidues are conserved. Thus, in φ29-like phages, pRNA may play a role in nucleotide binding/release by affecting the position or dyanmics of the gate motif.

### ADP release is promoted by a *trans*-acting arginine

The ATP-bound MD simulations described above show how individual subunit interfaces can bind and hydrolyze ATP. As revealed by SMFS studies, after all five ATPs have been hydrolyzed during the DNA translocating burst, the motor coordinates exchange of ADP for ATP in a sequential and interlaced manner^19^. To begin to understand the atomistic basis of these events, we simulated the structure of gp11 with ADP bound in each active site. The system was equilibrated with bound ADP and its equilibrium dynamics were simulated for at least 2.4 μs. Over this long sampling period, we were able to capture the initial steps of ADP-release. The α- and β-phosphates of ADP detached from the Walker A phosphate-binding motifs, while adensoine interactions were maintained. Importantly, release of the β-phosphate by the Walker A Lys32 was observed to be concomitant with interaction of the β-phosphate with the *trans*-acting Arg177 (**Fig. 3, Movie S1**). Thus, our simulations indicate that this arginine functions to promote nucleotide exchange, rather than as the canonical catalytic arginine finger. This observation provides atomistic explanation for recent SMFS observations that suggested a role for the analogous φ29 gp16 Arg146 in nucleotide exchange^23^.

**Figure 3:**
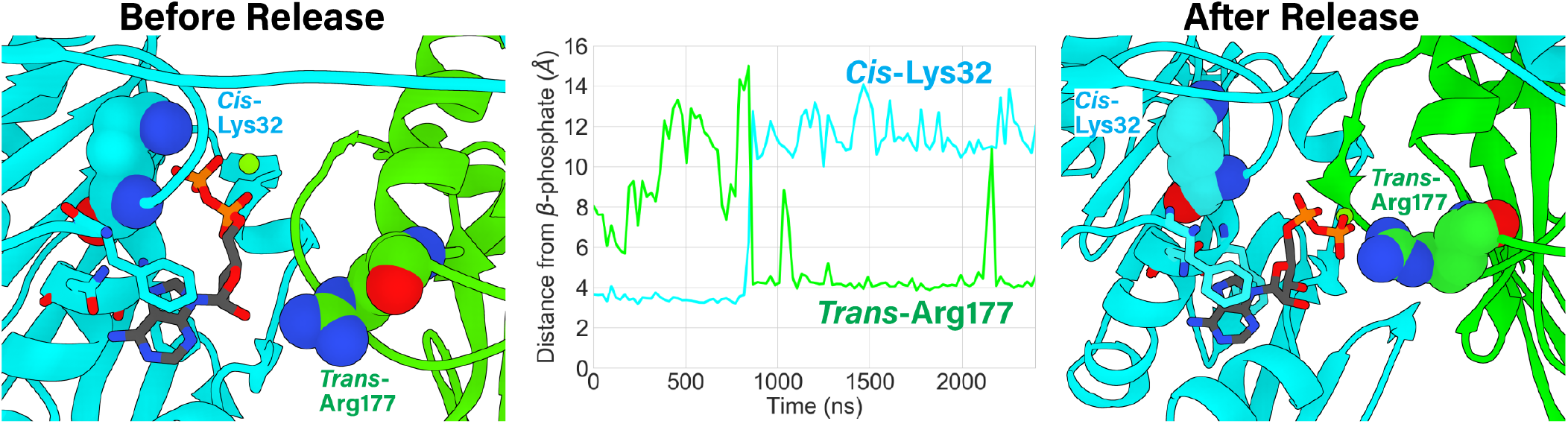
ADP-release is promoted by a trans-acting exchange residue. ADP unbinding is characterized by dissociation of the β-phosphate from the *cis*-acting Walker A Lys32 (interaction shown left panel) and concomitant association (time-evolved distance shwon middle panel) of the β-phosphate to *trans*-acting Arg177 (interaction shown right panel), which is implicated as being the exchange residue. Residues interacting with the adenosine (Phe8, Phe34, Asn35, shown unlabeled) largely maintain their interactions, acting as a pivot point to remove the phosphates from the binding pocket. The *cis*-acting Lys32 and *trans*acting Arg177 are shown as spheres and are labeled. The *cis*-acting enzyme is light blue, and the *trans*-acting enzyme is green.

After the phosphates were released from the Walker A motif, the Walker A backbone amide groups repositioned such that they could no longer bind the β-phosphate of ATP/ADP (**Movie S2**). This prevented ADP from accessing energetically favorable hydrogen bonding with the Walker A motif, reducing overall nucleotide affinity and likely precipitating complete dissociation of ADP from the active site. This phenomena has been characterized in myosin, where propensity of the Walker A motif to adopt a pose that is not receptive to β-phosphate hydrogen bonding is reported to be a predictor of ADP release rates^36^.

While our simulations were longer than typical MD simulations, they were not long enough to predict diffusion of ADP into solution. Nonetheless, we were interested in understanding further structural rearrangements that might occur upon complete dissociation of ADP. Thus, we performed additional simulations to probe the conformation and dynamics of a 4-ADP-bound, 1-apo subunit motor by removing the partially-unbound ADP described above from the system. We observe that the apo subunit’s Walker A motif backbone continued to rearrange, and finally positioned the Walker A Val30 in the “blocking” pose observed in the iodine crystal structure described above (**Movie S3**). Thus the simulations suggest that this pose may not be an artifact caused by iodine binding, but rather may be a mechanism that regulates ADP release and subsequent ATP binding.

### Positively-charged residues line the DNA pore

The crystallographic structures show that the pore through which DNA translocates is lined with several positively charged residues (**Fig. S4**). In the C-terminal domain, Lys332 and Lys336 point directly toward the channel lumen, and are thus well positioned to interact with DNA. A slight rotation of the C-terminal domain relative to the N-terminal domain would cause these side-chains to rotate out of the channel center and position Lys333, Lys334, and Lys366 in the lumen. Hence, both sets of residues may play a role in the packaging process, though in different stages of the mechanochemical cycle. This possibility is supported by the cryo-EM structure of φ29 particles imaged during packaging, which showed that residues approximately equivalent to Lys333, Lys334, and Lys366 interact with the substrate DNA during the dwell phase, presumably to prevent DNA slippage while the motor resets and exchanges ADP for ATP^11,18^. Hence, these residues likely function similarly in asccφ28.

The N-terminal (ATPase) domain also donates positively charged residues into the pore of the pentameric ring, namely Lys66, Lys92, Lys107, Arg110, and Arg128 (**Fig. 1**). While N-terminal domain-DNA interactions will be described in greater depth below, it is worth noting that these residues are well-conserved among other viral packaging ATPases, suggesting a conserved function (**Table S2**). Of note, asccφ28 Arg110 in is the only strictly conserved positively charged residue in all solved dsDNA viral packaging ATPases (**Table S2**). It was previously shown that the analogous Arg101 from the bacteriophage P74-26 packaging ATPase is absolutely necessary to bind substrate DNA^25,37^, suggesting a direct role in DNA translocation. The position of this residue in the interior of the pore supports this assignment.

### DNA-ATPase interactions depend on nucleotide-bound state

It is well known that the affinity for biopolymer substrates in ring ATPases depends on whether the ATPase active site is empty or if it has ATP or ADP bound. However, the structural basis for these changes in affinity is poorly understood. Thus, to assess the motor’s nucleotide-dependent affinity for substrate DNA, we systematically compared the DNA-binding residues from the ATP- and ADP-bound simulations described above (which included DNA in the pore). To complete the comparison, we performed equivalent 2.4 μs simulations of an apo pentamer with substrate DNA positioned in the pore. We observed that both the orientation of the positively-charged residues and the overall shape of the pore depend on nucleotide occupancy of the ATPase interfacial active sites.

Amongst the positively charged residues located in the lumen, we find that Arg110 is the most directly affected by nucleotide occupancy (**Fig. 4A**). In the all-apo simulation, all but one Arg110 lay flat along the interior of the pore, and do not interact strongly with DNA. When all five interfaces are bound with ATP, Arg110 is re-positioned into the pore as “prongs” that help grip DNA tightly. In the simulation, the planar arrangement of Arg110 prongs distort the helicity of DNA as the phosphate backbone attempts to interact with these positive charges (**Fig. 4B, left**). In the ADP-bound state, we find that Arg110 is closer to the apo position and do not protrude as prongs into the pore. These observations explain why an arginine at this position is so highly conserved and provide new insight into the general phenomenon of ATP-dependent increase in motor affinity for biopolymer substrates.

**Figure 4:**
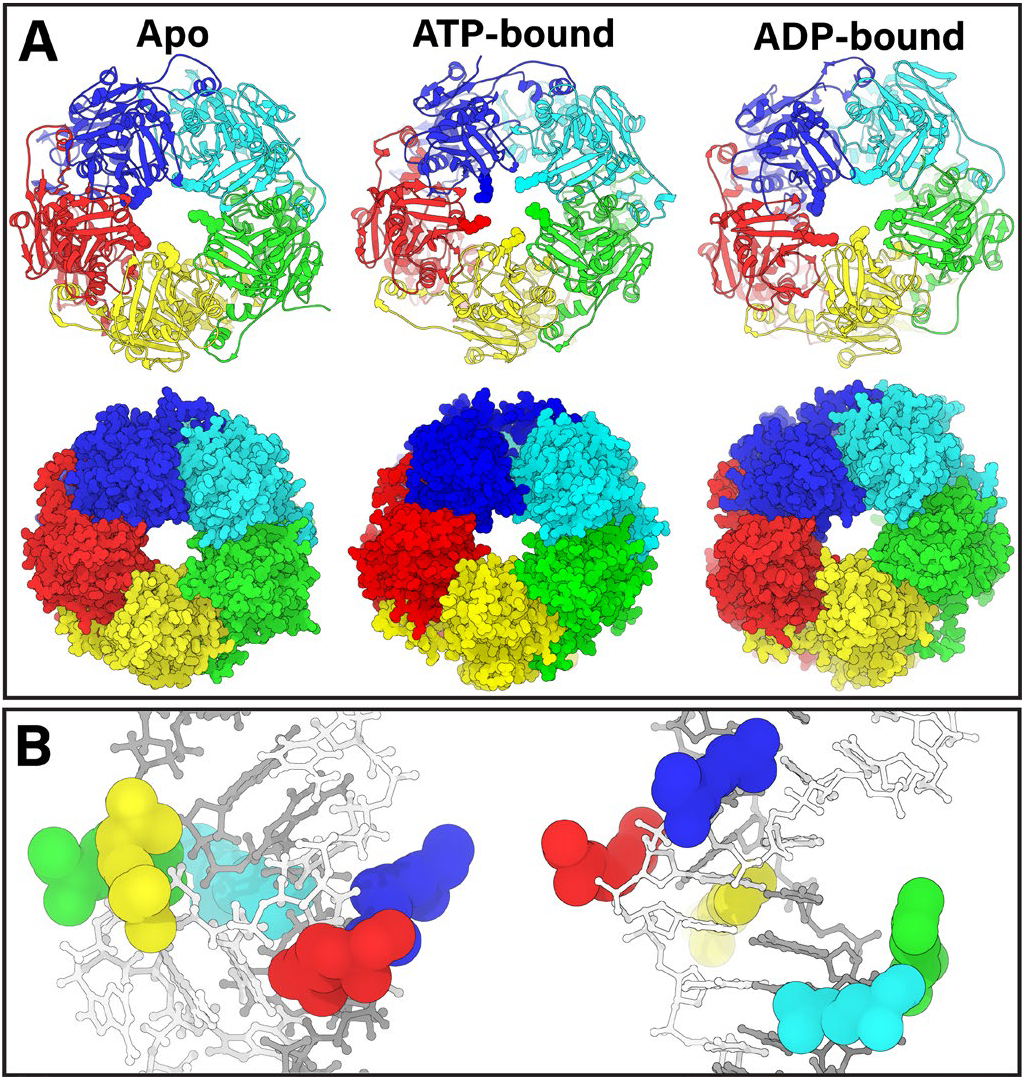
Pore geometry is modulated by nucleotide occupancy. **A.** Pentamer complex simulated in apo, ATP-bound, and ADP-bound states with substrate DNA (not shown for easier visualization) as Richardson diagrams (**top row**) and space-filling representations (**bottom row**). In the Richardson diagrams, Arg110 is shown as spheres to highlight its contribution towards DNA-gripping in the ATP-bound state. The pore is less constricted in the apo and ADP-bound states than the ATP-bound state, consistent with experimental data. **B.** Arg110 as predicted from the full-length subunit simulations are mostly a planar-ring and distort the DNA structure (**left**). Arg110 as predicted from ATPase-domain subunit simulations are donated following the helical pitch of DNA and do not distort the DNA structure (**right**).

### Lid subdomain flexibility provides molecular basis of force generation and motor reset

To quantitatively characterize the dynamics of the asccφ28 packaging ATPase obtained from our equilibrium MD simulations, we performed principal component analysis (PCA) coupled with root-mean-square fluctuation (RMSF) calculations on the alpha carbons of every residue in the gp11 ring in the different nucleotide bound states. RMSF provides us with a measure of how flexible a residue or motif is, while PCA allows us to understand whether flexibility is correlated as concerted motion along a specific direction or if the motion is essentially random (**Figs. 5 & S5**).

**Figure 5:**
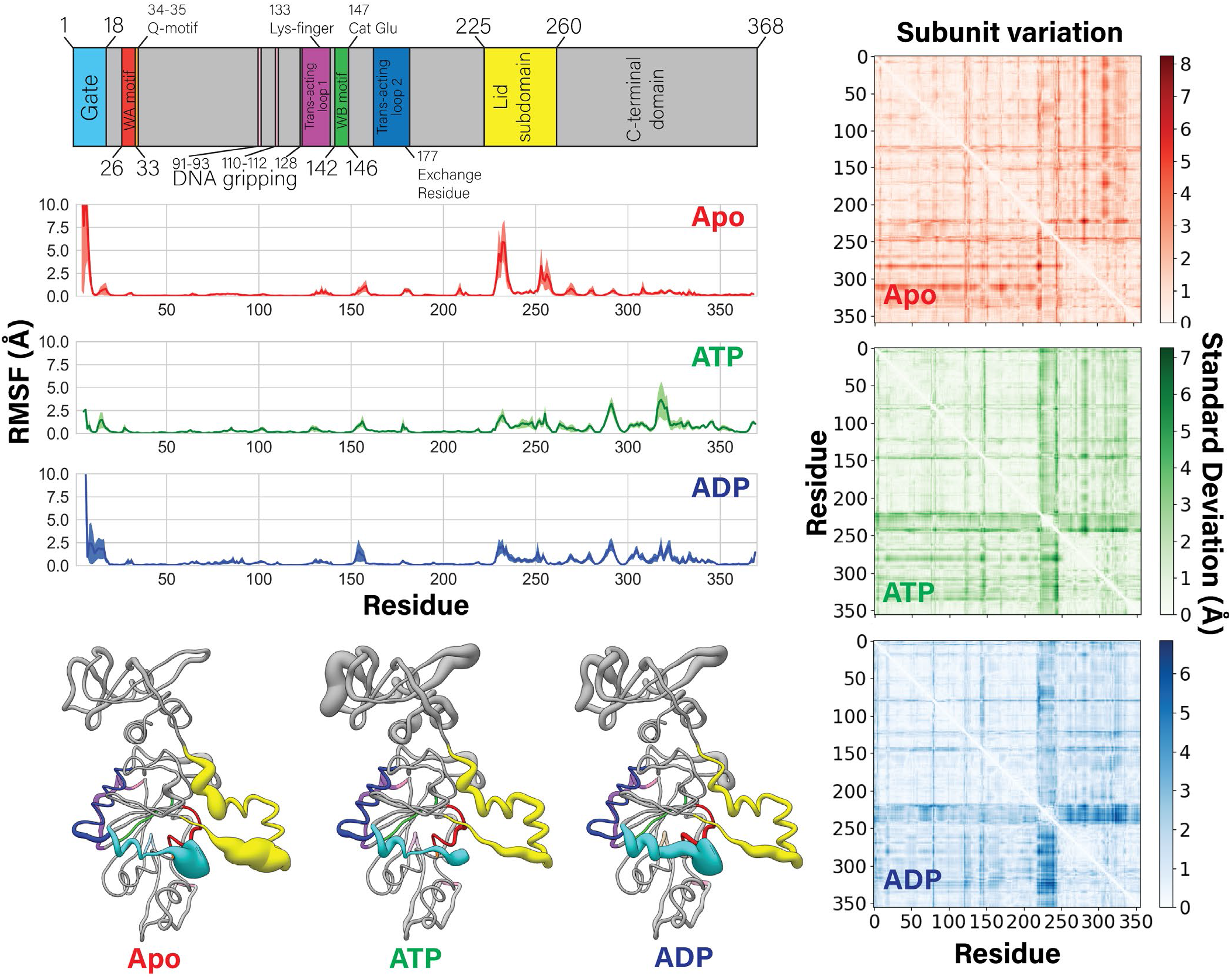
Flexibility of subunits is modulated by nucleotide occupancy. (**Left column**) Motif identity chart is reproduced at the top for easy reference. Averaged root-mean-square fluctuations (RMSF) of the ATPase alpha carbons over 2.4 microseconds of MD simulations are plotted. The standard error of the mean is shaded in each plot. Beneath the plots, the average RMSF value is coded to the radius of a worm-representation of the enzyme, which is color coded to the sequence. Thicker radius indicates higher flexibility. The apo state has a significantly more flexible lid subdomain than the ATP-or ADP-bound monomers. (**Right column**) Intra-subunit inter-residue distances are calculated in the apo, ATP-, and ADP-bound simulated states. The results from each of the five subunits are averaged. The standard deviations of the average are plotted as heatmaps, indicating structural variation across subunits within the pentamer. The ATP- and ADP-bound monomers have a band of high standard deviation in the lid subdomain (residues ~225-245), indicating that although each lid subdomain is rigid, they are in different poses. This band is muted in the apo state, indicating that the average positions of the lid subdomains are roughly equivalent, despite their flexibility. These observations correlate well with the asymmetry of the ATP- and ADP-bound pentamers, and the symmetry of the apo pentamer, given that the lid subdomain mediates most of the inter-subunit contacts.

Our analysis shows that the apo state is characterized by highly flexible N-terminal gate and lid subdomains. Flexibility of the lid subdomain can be attributed to the coils connecting the lid subdomain to the the N- and C-terminal domains. Thus, the helix-loop-helix motif of the lid subdomain moves as a rigid body, allowing it to maintain inter-subunit contacts with, and impart force on, its neighboring subunit.

Upon ATP binding, the simulations predict that both the N-terminal gate and the lid subdomains lose flexibility, and that the first principal component of motion is a rotation of the lid subdomain towards the ATPase active site (**Fig. S5**). Because the lid subdomain largely mediates inter-subunit contacts, lid subdomain rotation would pull two subunits closer together, enabling *trans*-acting catalysis; the mechanistic implications of this rotation are described below. In contrast, the ADP-bound state on average is characterized by a highly flexible N-terminal gate motif, but a rigid lid subdomain. However, the subunit wherein we observe ADP-release has a concerted rotation of the lid subdomain away from the ATPase active site (**Fig. S5**). This again suggests that ATP binding/ADP unbinding causes rotation of the lid subdomain. The dynamics described above in the pentamer simulations are echoed by similar dynamics observed in short-timescale simulations of a single subunit in the apo, ATP-bound, and ADP-bound states (**Movies S4, S5, S6**), which show that nucleotide binding rotates the lid subdomain over the ATPase active site.

To identify other regions of the protein that might be affected by lid subdomain rotation, we calculated intra-subunit residue-residue pairwise distances for all five subunits, and plotted the standard deviation of the average distance (**Fig. 5**). High standard deviation of average residue-residue distances indicates that the relative positions of residues vary across the five subunits; low standard deviation indicates structural homogeneity. The region of highest-standard deviation observed in the ATP- and ADP-bound simulated states corresponds to the lid subdomain. Likewise analysis of the three crystal structures shows that most of the variance is concentrated in the lid subdomain (**Fig. S6**). There is a second band of significant variation, which corresponds with the *trans*-acting lysine finger and its adjacent residues (residues 130-133) (**Fig. 5**, **S7**). Thus, not only does the lid subdomain directly contact *trans-*acting residues (**Fig. 1**), this analysis further indicates that the lid subdomain can modulate the positions of the key catalytic *trans*-acting residues. In summary, nucleotide-actuated rotation of a subunit’s lid subdomain can rearrange the motor’s overall quaternary structure and finely tune the position of *trans*-acting catalytic residues for appropriate function.

### Helix tracking in viral DNA packaging ATPases

We recently solved the structure of the φ29 packaging ATPase attached to the prohead and stalled during packaging^18^. This asymmetric cryo-EM reconstruction showed that the N-terminal domains of the ATPase ring adopt a helical conformation as each subunit tracks the substrate DNA. However, the crystal structure of asccφ28 gp11 ring reported here shows no such helicity in the N-or C-terminal domains. Likewise, the above MD simulations do not predict that the ATPase ring adopts a helical conformation. We suspect that we do not observe helicity in the crystal structures due to lack of substrate DNA imparting helicity. Lack of helicity in the MD simulations is likely attributable to interactions between the N- and C-terminal domains that result in a large activation energy barrier between extended and compact conformations. Thus, this transition may be kinetically infeasible to sample on the microsecond timescale of our simulations.

To remove this barrier, we simulated a pentamer ring composed of truncated monomers which contain only the N-terminal and lid (sub)domains (residues 1-260). Similar truncations in AAA+ systems have helped reveal biologically relevant helical conformations in cryo-EM structures^38,39^. MD simulations of the truncated ATPase assembly suggested that the motor can transition between planar and helical arrangements, such that each subunit interacts with the helical phosphate backbone of one strand of DNA (**Fig. S7**). In these simulations, rather than DNA distorting to allow interaction with Arg110 as in full-length simulations (**Fig. 4A**), the subunits in the ring translate to accomplish the same result without distorting B-form DNA (**Fig. 4B**). The helical arrangement of Arg110 also agrees with both the placement of an analogous DNA-gripping residue observed in the cryo-EM structure of φ29^18^, and SMFS experiments that showed the motor primarily tracks along a single strand of DNA^40^. To test if this propensity of ATPase domains to track DNA could be a general feature of viral packaging ATPases, we simulated pentamer assemblies of the P74-26 and D6E packaging ATPases constructed *in silico.* Again, our simulations predicted that these structures transition from the starting planar ring configuration to a helical configuration (**Fig. S8**). Furthermore, the simulated structures of the three systems (asccφ28, P74-26, and D6E) fit well into the cryo-EM reconstruction of the φ29 motor stalled with ATP analog (**Movies S8-10**).

## Discussion

An emerging feature of ring ATPases is that subunits in the ‘ring’ arrange themselves as a helix as they track their biopolymeric substrates^39,41-44^. Of particular relevance, a cryo-EM structure of φ29 stalled during packaging showed that its packaging motor adopts a helical pitch complementary to the double-stranded DNA substrate^18^. Based on this observation and the well-characterized behavior in SMFS studies^19,20,45,46^, it was proposed that φ29-like packaging motors transition between helical and planar states to translocate DNA. While the SMFS data identified the intermediate states that define the mechanochemical cycle and the cryo-EM structure documented the helical end state of the cycle, these data only tell half the story. The structures and simulations described here tell the other half of the story by documenting the structure of the planar end state and revealing the molecular basis of the conformational changes that drive the mechanochemical cycle.

### Helical-to-planar ratchet mechanism

Based on the results presented here, we propose a mechanism for viral DNA packaging where the motor ratchets between extended helical and compressed planar configurations to translocate DNA (**Fig. 6**). In this model, the transition from the helical to planar states drives DNA translocation during the burst phase, while the transition back from the planar to the helical state resets the motor during the dwell phase.

**Figure 6:**
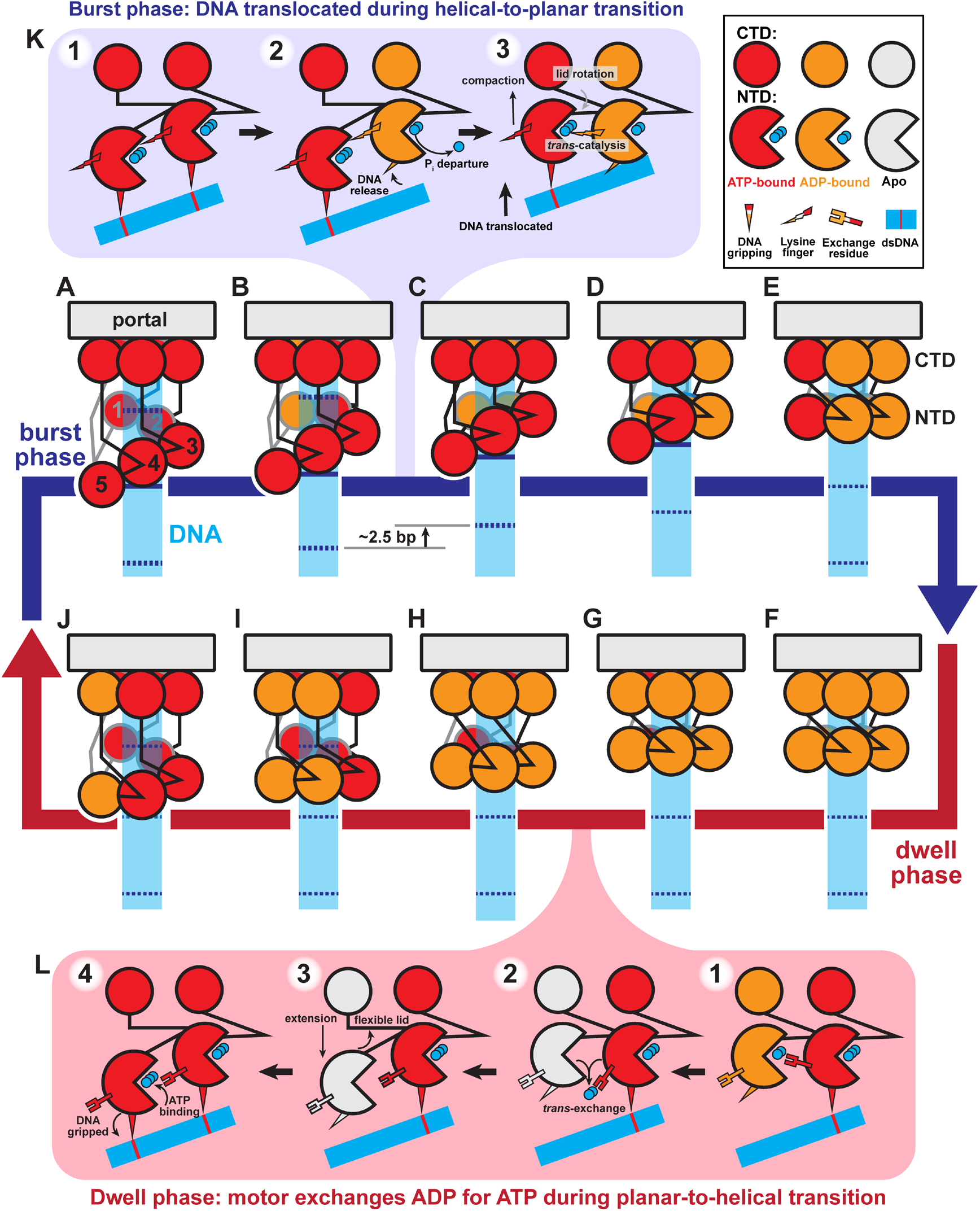
Helical-to-planar ratchet mechanism of DNA translocation. (**A-J**) Complete mechanochemical cycle. Subunits S1-S5 are labeled 1-5 in (**A**). During the burst phase (**A-E**), ATP-bound subunits (red) sequentially hydrolyze ATP. Hydrolysis in a subunit causes a pair of N-terminal domains (NTD) to become planar, translocating DNA. At the end of the burst, all subunits are ADP-bound (orange). During the dwell phase (**F-J**), ADP is sequentially exchanged for ATP, causing the planar NTD ring to return to the helical configuration. Helical-repeat-contacts of DNA (cyan) are marked by dashed lines. (**K**) Detailed schematic of the molecular events that coordinate the stepwise helical-to-planar transition. Initially, the two subunits’ NTD are ATP-bound, engaged with DNA, and helical. Hydrolysis and product release at the upper subunit relax its grip on DNA, allowing the other subunit’s lid subdomain to rotate, bringing the NTDs into a planar configuration, translocating DNA, and aligning the two subunits for the next hydrolysis event. (**L**) Detailed schematic of the molecular events that coordinate the stepwise planar-to-helical transition. After ADP-release is promoted by the *trans*-acting exchange residue, the now apo subunit (white) is flexible, and can track down the helix prior to ATP-binding, which causes the subunit to engage DNA, locking the subunit in place.

#### Mechanism of force generation

As depicted in **Fig. 6K**, ATP binding drives two competing effects. The predominant effect is increased affinity for DNA, resulting in the N-terminal domains adopting a helical configuration complementary to the helical phosphate backbone (**Figs. 4B and S7, Movies S8, S9, and S10**). The competing effect of ATP binding is lid subdomain rotation over the active site (**Fig. 5 and S5, Movies S4, S5, and S6**); because the lid subdomain is bound to a neighboring subunit (**Fig. 1**), this effect would drive subunits into the planar configuration if not for the interaction with the helical DNA. The tension between the two effects is resolved when a subunit hydrolyzes ATP and thus releases its grip on DNA; no longer constrained by its interaction with DNA, the lid subdomain of the adjacent ATP-bound neighbor can now rotate, bringing both subunits into a planar configuration. Since the ATP-bound subunit(s) maintains grip of DNA, this results in a discrete stepping of the DNA past the hydrolyzing subunit, through the ring and into the procapsid. Thus, resolution of the competing effects of ATP binding provides the basis of force generation and DNA translocation.

#### Burst

Our detailed description of the mechanochemical cycle starts when all five subunits are ATP-bound and therefore in the helical configuration (**Fig. 6A**). The subunit at the top of the helix (S1) hydrolyzes first and releases its grip on DNA, allowing its ATP-bound neighbor to rotate its lid subdomain and bring both subunits into a planar configuration (transition between **Fig. 6A** to **6B**). Since the remaining subunits (S2–S5) have yet to hydrolyze ATP, they continue to grip DNA such that ~2.5 bp of DNA are translocated into the procapsid. Upon planar alignment of the two N-terminal domains, the *trans*-acting lysine finger of the now-ADP-bound subunit is positioned to trigger hydrolysis in the adjacent ATP-bound subunit (**Fig. 6K; 2A**). Hydrolysis at this subunit initiates the next translocation step, as the hydrolyzing subunit releases grip of DNA, and lid subdomain of the ATP-bound adjacent subunit rotates, bringing both subunits into plane and translocating DNA as in the previous step (transition between **Fig. 6B** to **6C**). This pattern of hydrolysis at one subunit coordinating force generation of the adjacent subunit permutes around the ring until all N-terminal domains are in the planar configuration (**Fig. 6C-6E**). The result of one complete helical-to-planar transition is translocation of ~10 bp, or one helical turn, of DNA in four substeps. Thus, the helical-to-planar transition coordinated by sequential ATP hydrolysis constitutes the burst phase of the mechanochemical cycle.

#### Dwell

Once the motor is in the planar configuration, the final hydrolysis event would not translocate DNA (transition between **Fig. 6E** to **6F**). However, hydrolysis at the final subunit (S5) is needed to release DNA, such that the motor can step back down DNA during the reset. Additionally, single-molecule studies have shown that the fifth, non-translocating hydrolysis event plays a coordination role^19^. Thus, the final hydrolysis event likely coordinates the initiation of nucleotide exchange (**Fig. 6F**). The final subunit, whose N-terminal domain is now in plane with the N-terminal domain of the subunit that began the translocation burst (S1), donates its *trans*-acting arginine exchange residue to promote nucleotide exchange in the first subunit (transition between **Fig. 6F** to **6G; Fig. 3, Movie S1**). Because DNA was translocated one helical turn, by DNA’s translational symmetry, the subunit that began the burst (S1) is now positioned to reengage a DNA phosphate one helical turn below its last bound position. This highlights the importance of critical electrostatic contacts every helical turn of DNA observed in SMFS experiments^40^.

After ATP binds at the first subunit, it reengages DNA, locking its N-terminal domain in place. Then its *trans-*acting arginine exchange residue promotes nucleotide exchange in the adjacent subunit (S2) (transition between **Fig. 6G** to **6H**). Unlike the first subunit, the next subunit is misaligned with the helical DNA phosphate backbone. However, upon ADP release but before ATP binding, the subunit passes through the apo state where the lid subdomain is flexible (**Fig. 5**). This flexibility allows the subunit’s N-terminal domain to re-access the position where it is aligned with DNA (**Fig. 6L**). Because ATP binding and DNA binding are coupled (**Fig. 4**), this realignment would promote binding of ATP and tight binding of DNA, locking the N-terminal domain in place. As each subunit’s N-terminal domain samples the environment in the apo state, its Walker A motif may adopt the nucleotide-blocking conformation to prevent premature ATP binding (**Fig. S2, Movie S3**). This process permutes around the ring until the motor resets to the five ATP-bound, helical configuration that began the cycle (**Fig. 6H-A**). Thus, the planar-to-helical transition coordinated by sequential nucleotide exchange constitutes the dwell phase of the mechanochemical cycle.

#### Re-initiation of cycle

As described above, the first hydrolysis event occurs at the sheared interface, in the subunit closest to the capsid (S1 in **Fig. 6A**). The unique arrangement of this subunit’s lid subdomain as it reaches down to maintain contact across the sheared interface in the ATP-bound helical configuration positions a glutamine in its lid subdomain close to the γ-phosphate of ATP, as seen in Woodson et al.^18^. Hence, the glutamine is now poised to catalyze hydrolysis *in cis*. Thus, the motor re-initiates the burst phase when the ring adopts a fully helical configuration at the end of the planar-to-helical reset.

#### Implications of the mechanism

It has been reported that the φ29 packaging motor couples inorganic phosphate departure to DNA translocation^21^. However, this does not necessarily imply that phosphate departure provides the energy to drive the force-generating conformational changes in the motor. In our proposed model, ATP-binding creates tension between lid subdomain rotation promoting planarity and interactions with DNA promoting helicity. Hydrolysis and phosphate departure resolve this tension by releasing a subunit’s grip on DNA. This allows the ATP-binding energy stored within the neighboring strained lid subdomain to be converted into force exerted on the DNA. In this way, sequential phosphate departure serves as a trigger to initiate each stepwise movement of DNA.

Further, our mechanism also explains the physical basis of four translocation steps within the context of a motor with five subunits; only four steps are required to convert a pentameric helix into a planar ring. The last hydrolysis event (**Fig. 6E**), would not translocate DNA but coordinates nucleotide exchange, consistent with SMFS^19^. Additionally, the step size of the motor is set by the rise of the N-terminal domain helix rather than the periodicity of DNA and need not be coupled to an integer number of base pairs as has been demonstrated by SMFS^20^.

Perhaps the defining feature of φ29-like DNA packaging mechanochemistry is the biphasic burst-dwell cycle^19,20^ which would not result from inchworm or hand-over-hand mechanisms proposed for other helical ATPases^43,44^. In contrast, this key feature naturally emerges from our helical-to-planar ratchet mechanism. Compaction of the ring from the helical to planar configuration is the translocation burst (**Fig. 6A-E**), while extension of the ring from the planar to helical configuration is the nucleotide-exchange dwell (**Fig. 6F-J**). Thus, our model is consistent with and explains all data for φ29-like packaging motors, including recent SMFS measurements on the φ29 system packaging RNA and DNA/RNA hybrids that showed that the burst size readjusts to the periodicity of the substrate^45^. Hence, this type of mechanism may also explain the burst-dwell behavior of the polypeptide-translocating ClpX proteosome machinery^47^ and can serve as a general framework for any ring systems that are shown to exhibit burst-dwell dynamics.

## Supporting information

Methodology and Supplemental Information

## Author Contributions

Conceptualization, J.P., B.A.K, P.J., G.A., and M.C.M.; Methodology, J.P., E.D, M.A.W., B.A.K., P.J., G.A., and M.C.M; Investigation, J.P., E.D., and M.A.W.; Resources, P.J., G.A., and M.C.M.; Writing – Original Draft, J.P.; Writing – Review & Editing, J.P., B.A.K., P.J., G.A., M.C.M.; Visualization, J.P., B.A.K, P.J., G.A., and M.C.M.; Funding Acquisition, B.A.K., P.J., G.A., and M.C.M; Supervision, P.J., G.A., and M.C.M.

## Acknowledgments

This work was supported by the National institutes of Health grants GM122979 (to P.J.J. and M.C.M.), GM127365 (to M.C.M.), and GM118817 (to G.A.), and the National Science Foundation MCB1817338 (to B.A.K.). Computational resources for short equilibration simulations were provided by the NSF XSEDE Program ACI-1053575. Anton 2 computer time was provided by the Pittsburgh Supercomputing Center (PSC) through Grant R01GM116961 from the National Institutes of Health. The Anton 2 machine at PSC was generously made available by D.E. Shaw Research. We would also like to acknowledge the Sealy Center for Structural Biology and Molecular Biophysics (SCSB) for support of the UTMB structural and computational core facilities. The authors declare no conflict of interest.

## References

1. Erzberger, J. P. & Berger, J. M. Evolutionary relationships and structural mechanisms of AAA+ proteins. Annu. Rev. Biophys. Biomol. Struct. 35, 93–114 (2006).

2. Casjens, S. R. The DNA-packaging nanomotor of tailed bacteriophages. Nature reviews. Microbiology 9, 647–657 (2011).

3. Morais, M. C. The dsDNA packaging motor in bacteriophage ø29. in Viral Molecular Machines 511–547 (Springer, 2012).

4. Rao, V. B. & Feiss, M. Mechanisms of DNA Packaging by Large Double-Stranded DNA Viruses. Annual Review of Virology 2, 351–378 (2015).

5. Fuller, D. N. et al. Measurements of Single DNA Molecule Packaging Dynamics in Bacteriophage λ Reveal High Forces, High Motor Processivity, and Capsid Transformations. Journal of Molecular Biology 373, 1113–1122 (2007).

6. Fuller, D. N., Raymer, D. M., Kottadiel, V. I., Rao, V. B. & Smith, D. E. Single phage T4 DNA packaging motors exhibit large force generation, high velocity, and dynamic variability. Proceedings of the National Academy of Sciences 104, 16868–16873 (2007).

7. Pajak, J., Arya, G. & Smith, D. E. Biophysics of DNA Packaging. in Reference Module in Life Sciences B9780128096338209000 (Elsevier, 2019). doi: 10.1016/B978-0-12-809633-8.20966-7.

8. Smith, D. E. et al. The bacteriophage phi 29 portal motor can package DNA against a large internal force. Nature 413, 748–752 (2001).

9. Grimes, S., Jardine, P. J. & Anderson, D. Bacteriophage φ29 DNA packaging. (2002).

10. Ding, F. et al. Structure and assembly of the essential RNA ring component of a viral DNA packaging motor. Proceedings of the National Academy of Sciences 108, 7357–7362 (2011).

11. Mahler, B. P. et al. NMR structure of a vestigial nuclease provides insight into the evolution of functional transitions in viral dsDNA packaging motors. http://biorxiv.org/lookup/doi/10.1101/2020.07.06.188573 (2020) doi:10.1101/2020.07.06.188573.

12. Mao, H. et al. Structural and Molecular Basis for Coordination in a Viral DNA Packaging Motor. Cell Rep 14, 2017–2029 (2016).

13. Simpson, A. A. et al. Structure of the bacteriophage φ29 DNA packaging motor. Nature 408, 745–750 (2000).

14. Koti, J. S. et al. DNA packaging motor assembly intermediate of bacteriophage φ29. Journal of molecular biology 381, 1114–1132 (2008).

15. Mao, H. et al. Structural and Molecular Basis for Coordination in a Viral DNA Packaging Motor. Cell Reports 14, 2017–2029 (2016).

16. Morais, M. C. et al. Conservation of the capsid structure in tailed dsDNA bacteriophages: the pseudoatomic structure of φ29. Molecular cell 18, 149–159 (2005).

17. Morais, M. C. et al. Defining Molecular and Domain Boundaries in the Bacteriophage φ{symbol}29 DNA Packaging Motor. Structure 16, 1267–1274 (2008).

18. Woodson, M. et al. A viral genome packaging motor transitions between cyclic and helical symmetry to translocate dsDNA. http://biorxiv.org/lookup/doi/10.1101/2020.05.23.112524 (2020) doi:10.1101/2020.05.23.112524.

19. Chistol, G. et al. High Degree of Coordination and Division of Labor among Subunits in a Homomeric Ring ATPase. Cell 151, 1017–1028 (2012).

20. Moffitt, J. R. et al. Intersubunit coordination in a homomeric ring ATPase. Nature 457, 446–450 (2009).

21. Chemla, Y. R. et al. Mechanism of force generation of a viral DNA packaging motor. Cell 122, 683–692 (2005).

22. Reyes-Aldrete, E. et al. Biochemical and Biophysical Characterization of the dsDNA packaging motor from the Lactococcus lactis bacteriophage asccphi28. bioRxiv 2020.11.17.383745 (2020) doi:10.1101/2020.11.17.383745.

23. Tafoya, S. et al. Molecular switch-like regulation enables global subunit coordination in a viral ring ATPase. Proceedings of the National Academy of Sciences 115, 7961–7966 (2018).

24. Ogura, T., Whiteheart, S. W. & Wilkinson, A. J. Conserved arginine residues implicated in ATP hydrolysis, nucleotide-sensing, and inter-subunit interactions in AAA and AAA+ ATPases. Journal of Structural Biology 146, 106–112 (2004).

25. Hilbert, B. J. et al. Structure and mechanism of the ATPase that powers viral genome packaging. Proceedings of the National Academy of Sciences 112, E3792–E3799 (2015).

26. Cox, J. M., Abbott, S. N., Chitteni-Pattu, S., Inman, R. B. & Cox, M. M. Complementation of One RecA Protein Point Mutation by Another: EVIDENCE FOR TRANS CATALYSIS OF ATP HYDROLYSIS. J. Biol. Chem. 281, 12968–12975 (2006).

27. Mott, M. L., Erzberger, J. P., Coons, M. M. & Berger, J. M. Structural Synergy and Molecular Crosstalk between Bacterial Helicase Loaders and Replication Initiators. Cell 135, 623–634 (2008).

28. delToro, D. et al. Functional Dissection of a Viral DNA Packaging Machine’s Walker B Motif. Journal of Molecular Biology 431, 4455–4474 (2019).

29. Yang, R., Scavetta, R. & bao Chang, X. The hydroxyl group of S685 in Walker A motif and the carboxyl group of D792 in Walker B motif of NBD1 play a crucial role for multidrug resistance protein folding and function. Biochimica et Biophysica Acta-Biomembranes 1778, 454–465 (2008).

30. Davey, M. J., Jeruzalmi, D., Kuriyan, J. & O’Donnell, M. Motors and switches: AAA+ machines within the replisome. Nat Rev Mol Cell Biol 3, 826–835 (2002).

31. Sun, S., Kondabagil, K., Gentz, P. M., Rossmann, M. G. & Rao, V. B. The Structure of the ATPase that Powers DNA Packaging into Bacteriophage T4 Procapsids. Molecular Cell 25, 943–949 (2007).

32. Zhang, X. & Wigley, D. B. The ‘glutamate switch’ provides a link between ATPase activity and ligand binding in AAA+ proteins. Nature Structural and Molecular Biology 15, 1223–1227 (2008).

33. Tsay, J. M., Sippy, J., Feiss, M. & Smith, D. E. The Q motif of a viral packaging motor governs its force generation and communicates ATP recognition to DNA interaction. Proceedings of the National Academy of Sciences 106, 14355–14360 (2009).

34. Bhairosing-Kok, D. et al. Sharp kinking of a coiled-coil in MutS allows DNA binding and release. Nucleic Acids Research gkz649 (2019) doi:10.1093/nar/gkz649.

35. Zhan, X., Yan, C., Zhang, X., Lei, J. & Shi, Y. Structure of a human catalytic step I spliceosome. Science 359, 537–545 (2018).

36. Porter, J. R., Meller, A., Zimmerman, M. I., Greenberg, M. J. & Bowman, G. R. Conformational distributions of isolated myosin motor domains encode their mechanochemical properties. eLife 9, e55132 (2020).

37. Hilbert, B. J., Hayes, J. A., Stone, N. P., Xu, R. G. & Kelch, B. A. The large terminase DNA packaging motor grips DNA with its ATPase domain for cleavage by the flexible nuclease domain. Nucleic Acids Research 45, 3591–3605 (2017).

38. Han, H. et al. Structure of spastin bound to a glutamate-rich peptide implies a hand-over-hand mechanism of substrate translocation. J. Biol. Chem. 295, 435–443 (2020).

39. Zehr, E. A., Szyk, A., Szczesna, E. & Roll-Mecak, A. Katanin Grips the β-Tubulin Tail through an Electropositive Double Spiral to Sever Microtubules. Developmental Cell 52, 118–131.e6 (2020).

40. Aathavan, K. et al. Substrate interactions and promiscuity in a viral DNA packaging motor. Nature 461, 669–673 (2009).

41. De la Peña, A. H., Goodall, E. A., Gates, S. N., Lander, G. C. & Martin, A. Substrate-engaged 26S proteasome structures reveal mechanisms for ATP-hydrolysis–driven translocation. Science 362, (2018).

42. Enemark, E. J. & Joshua-Tor, L. Mechanism of DNA translocation in a replicative hexameric helicase. Nature 442, 270–275 (2006).

43. Jean, N. L., Rutherford, T. J. & Löwe, J. FtsK in motion reveals its mechanism for doubl-estranded DNA translocation. Proc Natl Acad Sci USA 117, 14202–14208 (2020).

44. Monroe, N., Han, H., Shen, P. S., Sundquist, W. I. & Hill, C. P. Structural basis of protein translocation by the Vps4-Vta1 AAA ATPase. eLife 6, 1–22 (2017).

45. Castillo, J. P., Tong, A., Tafoya, S., Jardine, P. J. & Bustamante, C. A DNA translocase operates by cycling between planar and lock-washer structures. http://biorxiv.org/lookup/doi/10.1101/2020.05.22.101154 (2020) doi:10.1101/2020.05.22.101154.

46. Liu, S. et al. A viral packaging motor varies its DNA rotation and step size to preserve subunit coordination as the capsid fills. Cell 157, 702–713 (2014).

47. Maillard, R. A. et al. ClpX(P) Generates Mechanical Force to Unfold and Translocate Its Protein Substrates. Cell 145, 459–469 (2011).

