## Supplementary material for "Atomistic Basis of Force Generation, Translocation, and Coordination in a Viral Genome Packaging Motor": Methodology and Supplemental Information

### Structure determination

Crystallization. Protein in buffer containing 50 mM sodium phosphate, pH 8.1, 400 mM sodium chloride, and 1 mM dithiothreitol was concentrated via filtration to 4.3 mg/ml. Several commercial sparse matrix screens were used to determine initial crystallization conditions, and initial crystals were obtained from the Wizard Classic screen (Rigaku) in 2 M ammonium sulfate, 0.1 M sodium citrate, pH 5.5 and Salt RX screen (Hampton Research) in 0.7 M citrate, 0.1 M Tris, pH 8.5. Gp11 has a predicted pI of 6.44, so would be positively or negatively charged, respectively, in the two conditions. Each condition served as the starting point for several optimization steps, and crystals exceeding 100μm were eventually grown in 24-well VDX trays by hanging-drop diffusion over 1000 μL of well solution. A rhombohedral crystal (Crystal Form A) was grown over wells containing 1.5-1.8 M ammonium sulfate, 0.1 M citrate, pH 5.3-5.7. The drop consisted of 1μl protein at 4.3 mg/ml mixed with 1μl well solution containing 1.6 M ammonium sulfate, 0.1 M trisodium citrate, pH 5.7. The pH of trisodium citrate buffers was adjusted with hydrochloric acid. Crystals appeared after 2-4 days. Octahedral crystals (Crystal Form B) were grown from basic conditions containing 1.0 M trisodium citrate, 0.1 M Tris pH 8.3. The final pH of this well solution was measured at 8.9. Protein concentration was 3.1 mg/ml, and 1.5 μL of protein was combined with an equal volume of well solution. Both solutions were pre-chilled, and the tray was set up and incubated at 277 K. Well-formed crystals typically took more than a month to grow.

X-ray Crystallographic data collection. Crystals of asccφ28 gp11 were soaked in mother liquor with 20% glycerol added as a cryoprotectant prior to flash-freezing. The x-ray diffraction data were collected at the Advanced Photon Source (APS) Life Science Collaborative Access Team (LS-CAT) beamlines (**Table S1**). The data were indexed, scaled, and merged using **HKL2000** (**Table S1**) (Minor et al., 2006).

Structure solution and model building. The $P3_221$ structure, grown in 1.6 M ammonium sulphate and 0.1 M trisodium citrate, pH 5.7, was solved to 3.3 Å via SAD phasing using the programs **Shelx** (Sheldrick, 2008) and **HKL2MAP** (Pape and Schneider, 2004). The data were from a SeMet derivatized crystal. An initial model was built using a **Parrot** (Cowtan, 2010) NCS-averaged map in **Bucaneer** (Cowtan, 2006), both from **CCP4** program suite (Winn et al., 2011). The final structure was refined to 2.9 Å using **Phenix** (Adams et al., 2010) and built in **COOT** (Emsley et al., 2010). The $P4_32_12$ native structure, grown in 1.0 M trisodium citrate, 0.1 M tris (hydroxymethyl) aminomethane (Tris), pH 8.3, was solved using molecular replacement in Phaser (McCoy et al., 2007), using the pentaric ring from the $P3_221$ cell as search model. The structure was refined using **Phenix** (Adams et al., 2010). Model-building was performed using **COOT** (Emsley et al., 2010). The $P4_32_12$ Iodide-soaked structure was solved via direct phasing using the previous $P4_32_12$ structure described above as a starting model. The structure was refined using **Phenix** (Adams et al., 2010). Model-building was performed using **COOT** (Emsley et al., 2010). Final refinement statistics for all structures are summarized in (**Table S1**).

### Molecular dynamics simulations

Structure preparation. All molecular dynamics (MD) simulations of the pentameric ATPase ring of bacteriophage asccφ28 were started from the crystal structures reported herein. Missing side chains were added to the structures by using the Dunbrack rotamer library (Dunbrack Jr, 2002). The "ATP-bound" pentamer was prepared by docking ATP into the binding pockets based on ATP-bound structures of homologous viral ATPases. Docking included ATP, $Mg^{2+}$, and a water molecule situated between $Mg^{2+}$ and the Walker B Asp, which is crucial to maintain the canonical hydrogen bond coordination within the binding pocket. The "ADP-bound" pentamer was prepared by removing the γ-phosphate from the initial coordinates of the ATP-bound subunits. The "4-ADP bound, 1-apo mixed occupancy" pentamer was prepared by removing the unbinding ADP molecule from the final frame of the 5-ADP bound simulation. A 30 bp-long B-form double-stranded DNA molecule was generated by using the 3DNA webserver (Li et al., 2019) and manually placed into the pore of the ATPase ring, which is large enough to accommodate B-form DNA with minimal steric clashes. The "ATPase-domain only" pentamer was generated by truncating the C-terminal domains (residues 261 onwards) from the crystal structure, and a shorter 25 bp-long double-stranded DNA molecule was placed in the pore. Monomer simulations were started from one arbitrarily chosen subunit of the pentamer complex.

The pentamer of the P74-26 ATPase was constructed from the solved crystal structure of the ATPase domain P74-26 (PDB: 4ZNL) by using the M-ZDOCK protein-protein docking software (Pierce et al., 2005), imposing five-fold symmetry. The structure produced matched the reported prediction (Hilbert et al., 2015), which was likewise generated with M-ZDOCK. ATP-bound structures were generated by placing $Mg^{2+}$-ATP in the binding pocket according to the solved crystal structure of monomeric P74-26 ATPase containing bound ADP-$BeF_3$ in the active site. As done for the asccφ28 pentamer, a double-stranded B-DNA molecule was generated with 3DNA and manually placed in the pore of the structure. Due to the larger size of the P74-26 ATPase, the molecule length was extended to 35 bp. After equilibration, the identified arginine finger was poised to catalyze hydrolysis in one and only one active site. To probe the effects of subsequent hydrolysis, ATP was removed from this active site, and the system equilibrated again. During this equilibration, the system transitioned from a planar ring to a helical arrangement, shearing at the single apo interface (**Fig. S9**).

Lastly, the D6E ATPase pentamer was constructed by superimposing a monomer structure (PDB: 5OE8) (Xu et al., 2017) which had undergone 2.4 microseconds of equilibrium MD simulation onto the asccφ28 crystal structure reported here. During the simulation, the monomer's lid subdomain extends away from the ATPase domain (**Movie S7**), allowing for ample inter-subunit contacts similar to the asccφ28 crystal structure reported here. The C-terminal domain was truncated to prevent inter-subunit steric clash. A 25 bp-long double-stranded DNA molecule generated using 3DNA was manually placed into the pore of the structure.

*Pentamer simulations.* All-atom, explicit-solvent MD simulations of all pentamers were carried out using the Anton 2 simulation package and supercomputer (Shaw et al., 2014). However, the equilibration runs, including structure preparation and energy minimization, were all performed using the AMBER18 simulation package with GPU optimization (Salomon-Ferrer et al., 2013), as all systems must be equilibrated before production runs can be performed on Anton 2. The equilibration and production simulations both used the AMBER99SB-ILDN force field to describe protein interactions (Lindorff-Larsen et al., 2010), and the bsc1 AMBER parameter update to describe DNA interactions (Ivani et al., 2015). ATP and ADP parameters are taken from the AMBER parameter database (Meagher et al., 2003).

For equilibration using AMBER18 package, all systems were centered in a periodic cubic box of TIP3P water with 14 Å minimum padding. The systems were charge-neutralized with counterions and additional salt was added to reach 150 mM concentration. The systems were energy minimized using a combination of steepest descent and conjugate gradient, and then slowly heated in the NVT ensemble from 100 K to 310 K over 100 ps using the Langevin thermostat. Bonds connecting heavy atoms to hydrogen were constrained through the SHAKE algorithm, and a 2-fs integration time step was used to propagate the equations of motion. Subsequently, each system was subjected to short simulation in the NPT ensemble, held at 310 K via the Langevin thermostat and 1 bar with the Monte Carlo barostat to equilibrate the density. Finally, each system was simulated for at least 2.4 microseconds on the Anton 2 supercomputer. These production runs were in the NPT ensemble held at 310 K and 1 bar using the Nose-Hoover thermostat and the MTK barostat and a 2.5 fs integration time step to propagate the equations of motion.

*Characterizing subunit motions and flexibility*. Principal components (PCs) and root-mean-square fluctuations (RMSF) were calculated for each subunit individually by aligning the trajectories to minimize root-mean-square deviation (RMSD) of that single subunit. Thus, the PCs and RMSF represent internal degrees of freedom of a subunit and do not account for motion of a subunit relative to the rest of the pentamer. These quantities were calculated on a per-residue basis using the α-carbons in each residue excluding the side chain. Thus, the obtained values reflect backbone motion and not rotamer states. PCs and RMSF were calculated using ProDy software as interfaced with VMD. RMSF curves reported in main text **Fig. 5** were calculated by averaging the per-residue RMSF of all five subunits within a single trajectory, and the uncertainty reported is the standard error of the mean.

Subunit variation (main text **Fig. 5** and **Fig. S7**) was determined by calculating the residue-residue pairwise distances within a single subunit for each subunit of a given pentamer structure. Then the five values were averaged, and the standard deviation of this average is plotted as heat maps in main text **Fig. 5**. High standard deviation in the average residue-residue pairwise distance indicates structural variability across the five subunits.

*Monomer simulations.* To better understand the dynamics of an individual subunit within the pentamer, we also performed 100 ns long all-atom explicit MD simulations of a monomer in the apo, ATP-bound, and ADP-bound states. These short timescale monomer simulations were performed exclusively in the AMBER18 simulation package with GPU acceleration. We performed simulations using both the AMBER99SB-ILDN/TIP3P and the AMBER ff19SB/OPC protein/water force field combinations. The reason for considering two separate force field combinations is provided below. These systems were centered in a periodic truncated-octahedron box of water to reduce the total number of particles and save computation time. The systems were equilibrated using the same procedure described above for pentamers. 100-ns production runs were performed in triplicate for both set of force fields in the NPT statistical ensemble, held at 310 K and 1 bar by the Langevin thermostat and Monte Carlo barostat.

The AMBER99SB-ILDN/TIP3P simulations allowed for direct comparison to the pentamer simulations performed on Anton 2 which used the same pairing of force fields. However, it has been shown that the TIP3P water model underpredicts the strength of protein-water interactions and promotes compact secondary structure formation. We were primarily interested in the dynamics of the lid subdomain, which is extended away from the ATPase domain via a flexible linker. In the pentamer structures, the lid subdomain forms extensive contacts with neighboring subunits, which may help stabilize it in its extended state. In the monomer simulations these protein-protein contacts are replaced by protein-water contacts. Thus, we also considered the new AMBER ff19SB/OPC force field pairing (Izadi et al., 2014; Tian et al., 2020), which strengthens the protein-water interaction and helps stabilize extended states. In the end, both force field pairings predicted similar conformations and dynamics of the lid subdomain on the timescale we considered.

SAMSON (https://www.samson-connect.net) was used to create main text **Fig. 1-4**. UCSF Chimera (Pettersen et al., 2004) was used to visualize main text **Fig. 5-6**.


Adams, P.D., Afonine, P.V., Bunkóczi, G., Chen, V.B., Davis, I.W., Echols, N., Headd, J.J., Hung, L.-W., Kapral, G.J., Grosse-Kunstleve, R.W., et al. (2010). PHENIX: a comprehensive Python-based system for macromolecular structure solution. Acta Crystallographica Section D: Biological Crystallography *66*, 213–221.

Cowtan, K. (2006). The Buccaneer software for automated model building. 1. Tracing protein chains. Acta Crystallographica Section D: Biological Crystallography *62*, 1002–1011.

Cowtan, K. (2010). Recent developments in classical density modification. Acta Crystallographica Section D: Biological Crystallography *66*, 470–478.

Dunbrack Jr, R.L. (2002). Rotamer libraries in the 21st century. Current Opinion in Structural Biology *12*, 431–440.

Emsley, P., Lohkamp, B., Scott, W.G., and Cowtan, K. (2010). Features and development of Coot. Acta Crystallographica Section D: Biological Crystallography *66*, 486–501.

Hilbert, B.J., Hayes, J.A., Stone, N.P., Duffy, C.M., Sankaran, B., and Kelch, B.A. (2015). Structure and mechanism of the ATPase that powers viral genome packaging. Proceedings of the National Academy of Sciences *112*, E3792–E3799.

Ivani, I., Dans, P.D., Noy, A., Pérez, A., Faustino, I., Hospital, A., Walther, J., Andrio, P., Goñi, R., Balaceanu, A., et al. (2015). Parmbsc1: A refined force field for DNA simulations. Nature Methods *13*, 55–58.

Izadi, S., Anandakrishnan, R., and Onufriev, A.V. (2014). Building Water Models: A Different Approach. J. Phys. Chem. Lett. *5*, 3863–3871.

Li, S., Olson, W.K., and Lu, X.-J. (2019). Web 3DNA 2.0 for the analysis, visualization, and modeling of 3D nucleic acid structures. Nucleic Acids Research *47*, W26–W34.

Lindorff-Larsen, K., Piana, S., Palmo, K., Maragakis, P., Klepeis, J.L., Dror, R.O., and Shaw, D.E. (2010). Improved side-chain torsion potentials for the Amber ff99SB protein force field. Proteins: Structure, Function and Bioinformatics *78*, 1950–1958.

McCoy, A.J., Grosse-Kunstleve, R.W., Adams, P.D., Winn, M.D., Storoni, L.C., and Read, R.J. (2007). Phaser crystallographic software. Journal of Applied Crystallography *40*, 658–674.

Meagher, K.L., Redman, L.T., and Carlson, H.A. (2003). Development of polyphosphate parameters for use with the AMBER force field. Journal of Computational Chemistry *24*, 1016–1025.

Minor, W., Cymborowski, M., Otwinowski, Z., and Chruszcz, M. (2006). HKL-3000: the integration of data reduction and structure solution–from diffraction images to an initial model in minutes. Acta Crystallographica Section D: Biological Crystallography *62*, 859–866.

Pape, T., and Schneider, T.R. (2004). HKL2MAP: a graphical user interface for macromolecular phasing with SHELX programs. Journal of Applied Crystallography *37*, 843–844.

Pettersen, E.F., Goddard, T.D., Huang, C.C., Couch, G.S., Greenblatt, D.M., Meng, E.C., and Ferrin, T.E. (2004). UCSF Chimera—a visualization system for exploratory research and analysis. Journal of Computational Chemistry *25*, 1605–1612.

Pierce, B., Tong, W., and Weng, Z. (2005). M-ZDOCK: A grid-based approach for Cnsymmetric multimer docking. Bioinformatics *21*, 1472–1478.

Salomon-Ferrer, R., Götz, A.W., Poole, D., Le Grand, S., and Walker, R.C. (2013). Routine Microsecond Molecular Dynamics Simulations with AMBER on GPUs. 2. Explicit Solvent Particle Mesh Ewald. J. Chem. Theory Comput. *9*, 3878–3888.

Shaw, D.E., Grossman, J.P., Bank, J.A., Batson, B., Butts, J.A., Chao, J.C., Deneroff, M.M., Dror, R.O., Even, A., Fenton, C.H., et al. (2014). Anton 2: Raising the Bar for Performance and Programmability in a Special-Purpose Molecular Dynamics Supercomputer. In SC14: International Conference for High Performance Computing, Networking, Storage and Analysis, (New Orleans, LA, USA: IEEE), pp. 41–53.

Sheldrick, G.M. (2008). A short history of SHELX. Acta Crystallographica Section A: Foundations of Crystallography *64*, 112–122.

Tian, C., Kasavajhala, K., Belfon, K.A.A., Raguette, L., Huang, H., Migues, A.N., Bickel, J., Wang, Y., Pincay, J., Wu, Q., et al. (2020). ff19SB: Amino-Acid-Specific Protein Backbone Parameters Trained against Quantum Mechanics Energy Surfaces in Solution. J. Chem. Theory Comput. *16*, 528–552.

Winn, M.D., Ballard, C.C., Cowtan, K.D., Dodson, E.J., Emsley, P., Evans, P.R., Keegan, R.M., Krissinel, E.B., Leslie, A.G., McCoy, A., et al. (2011). Overview of the CCP4 suite and current developments. Acta Crystallographica Section D: Biological Crystallography *67*, 235–242.

Xu, R.-G., Jenkins, H.T., Antson, A.A., and Greive, S.J. (2017). Structure of the large terminase from a hyperthermophilic virus reveals a unique mechanism for oligomerization and ATP hydrolysis. Nucleic Acids Research *45*, 13029–13042.

| Crystal | Se-Met | NaI | Native |
|---|---|---|---|
| PDB-ID | TBD | TBD | TBD |
| Source | 21-ID-F | 21-ID-G | 21-ID-F |
| Wavelength (Å) | 0.97872 | 0.97857 | 0.97872 |
| Detector | Rayonix MX300 | Rayonix MX300 | Rayonix MX300 |
| Spacegroup | $P3_221$ | $P4_32_12$ | $P4_32_12$ |
| Resolution (highest shell) | 2.8 (2.85-2.80) | 2.9 (2.95-2.90) | 2.9 (2.95-2.90) |
| **Cell Dimensions** | | | |
| a (Å) | 135.0 | 112.2 | 110.9 |
| b (Å) | 135.0 | 112.2 | 110.9 |
| c (Å) | 276.7 | 354.2 | 351.8 |
| A, β, γ (°) | 90 90 120 | 90 90 90 | 90 90 90 |
| No Frames | 720 | 234 | 240 |
| Osc Range (°) | 1 | 0.5 | 0.5 |
| No. Reflections | 5,302,794 | 2,924,141 | 3,203,867 |
| No. Merged Reflections (Anom) | 43,588 (62,234) | 42,759 (78,494) | 49,847 |
| Rpim | 0.028 (1.4) | 0.04.3 (1.1) | 0.05.7 (1.0) |
| CC1/2 | NA (0.52) | 0.99 (0.21) | 0.98 (0.30) |
| I/s | 16.8 (0.5) | 16.3 (0.6) | 11.0 (0.6) |
| Completeness (%) | 85 (31) | 99.7 (97) | 98.9 (90) |
| Redundancy | 43 (12) | 16.4 (7.5) | 10.9 (4.8) |
| **Refinement** | **Se-Met** | **NaI** | **Native** |
| **Phasing** | SAD | MR | MR |
| Resolution (highest shell) | 2.9 (2.94-2.90) | 2.9 (2.93-2.90) | 3.0 (3.03-3.00) |
| No. Unique Reflections | 43,583 | 42,759 | 38,744 |
| Rw (%) | 24.12 | 23.46 | 23.80 |
| Rf (%) | 27.41 | 28.37 | 27.31 |
| Bond (Angle) RMS (Å)(°) | 0.004 (0.6) | 0.004 (0.7) | 0.004 (0.9) |
| No. Molecules in a.u. | 5 | 5 | 5 |
| Non-hydrgon protein atoms avg. B (N) | 93 (15,401) | 58 (15,123) | 56 (15,065) |
| Non-hydrogen solvent atoms avg. B (N) | NA (0) | 81 (152) | 41 (31) |
| **Ramachandran** | | | |
| Favoured (%) | 97.80 | 97.36 | 97.22 |
| Allowed (%) | 2.20 | 2.48 | 2.73 |
| Outliers (%) | 0.00 | 0.17 | 0.06 |

**Table S1: Crystallographic Data and Refinement Statistics**

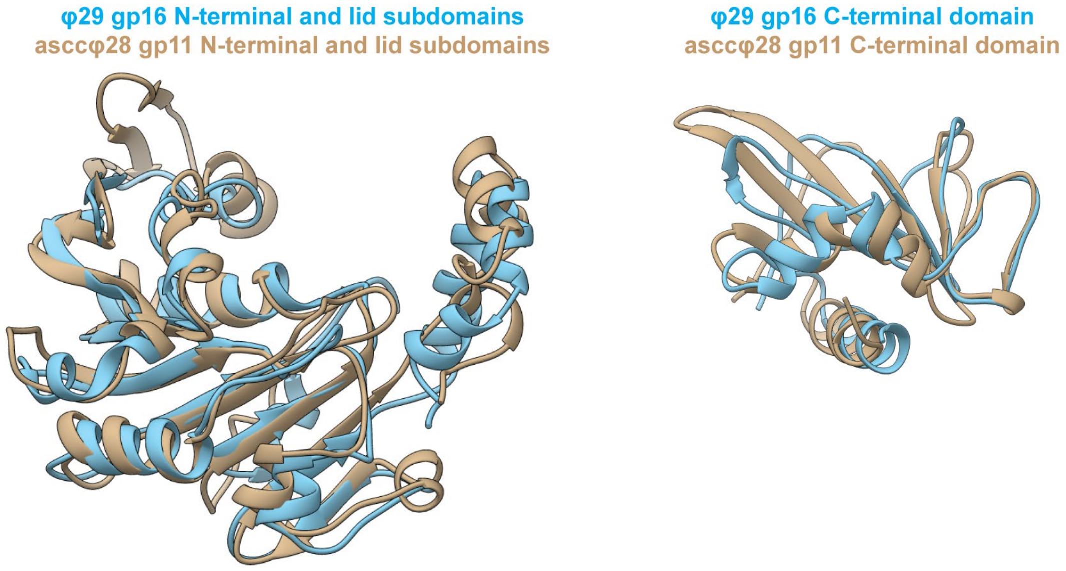


**Figure S1:** Superimpositions of the φ29 gp16 and asccφ28 gp11 domains show nearly identical structures, as expected based on their 45% sequence similarity.

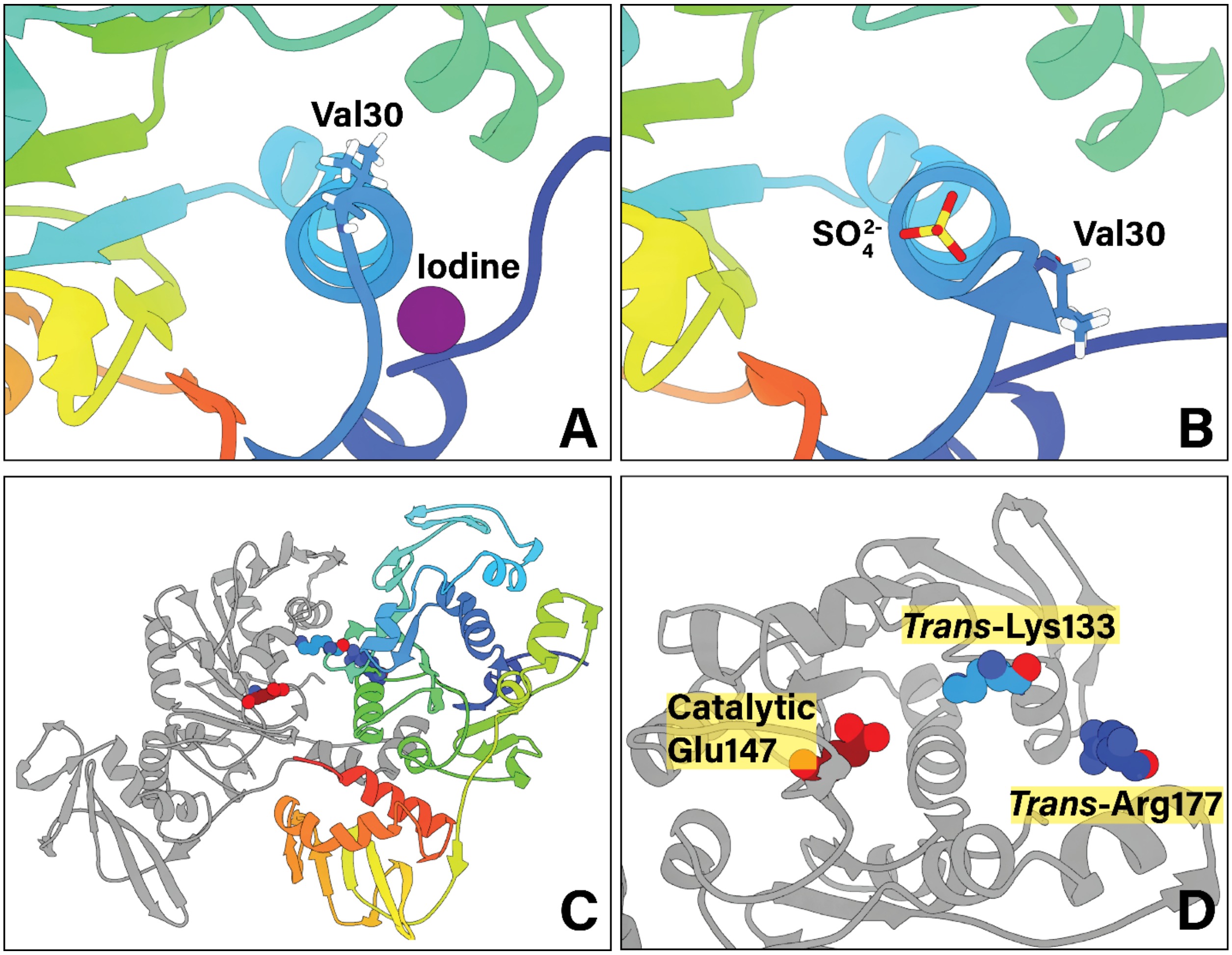


**Figure S2: The active site observed in the crystal structure.** Two crystals show that the Walker A motif varies helical content. Extra helical content (**A**) places Val30 in the binding pocket occluding nucleotide occupancy, and less helical content (**B**) points Val30 away from the binding pocket. This allows a sulfate ion to mediate interactions in the P-loop normally mediated by the β-phosphate of ATP/ADP. (**C**) A zoomed-out view of two neighboring subunits, shown in gray and rainbow. (**D**) Zoom in on the active site shows that the *trans*-acting Lys133 is considerably closer to the *cis*-acting catalytic Glu147 than *trans*-acting Arg177 is; from this alone it can be inferred that the lysine acts as a lysine finger, as it is better situated to interact with the γ-phosphate of ATP near the catalytic Glu147*.*

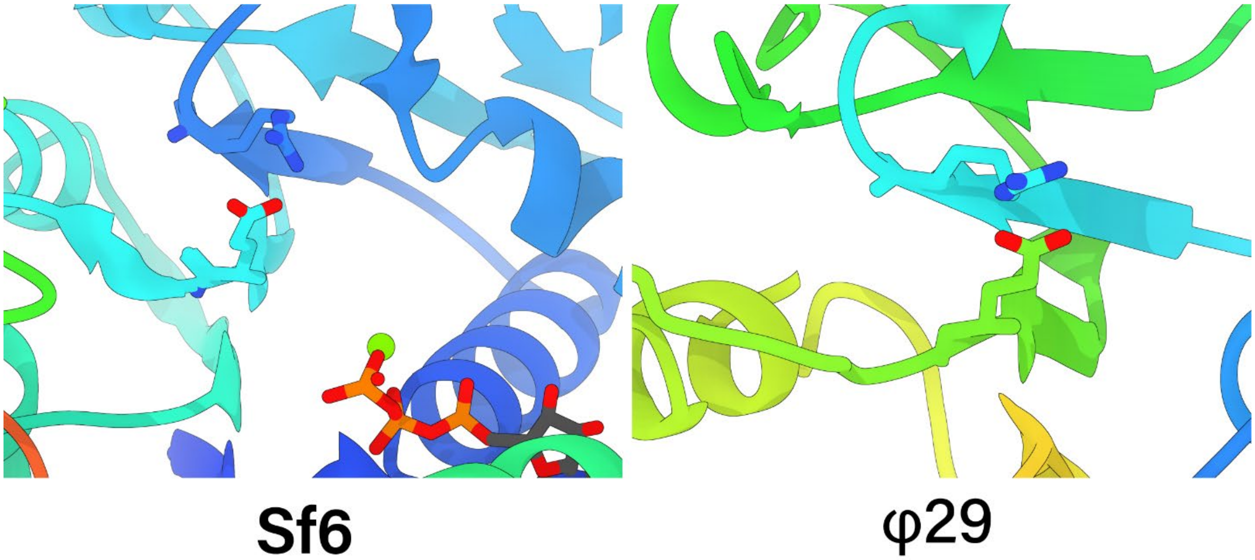


**Figure S3: The catalytic glutamate is observed to be inactive in the Sf6 and φ29 crystal structures.** The Sf6 and φ29 packaging ATPases are shown as Richardson diagrams, and zoomed into the ATPase active site. The catalytic glutamate is shown as sticks, pointing its carboxylate away from ATP and towards a conserved arginine residue also shown as sticks, reminiscent of the glutamate switch mechanism found in other AAA+ enzymes.

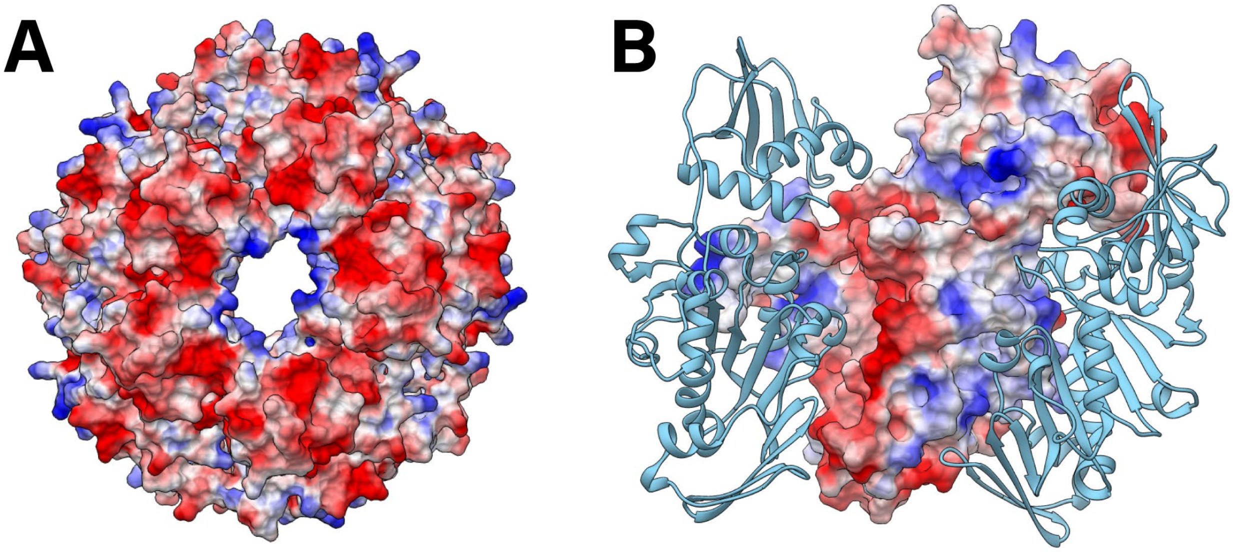


**Figure S4: Electrostatics of the pore.** (**A**) An end-on view of the gp11 pentamer crystal structure shows that the interior of the pore is lined with positive electrostatic surface potential. (**B**) A cut away view of the pentamer shows a positive band of electrostatic potential runs along the length of the monomer, creating a DNA-gripping strip. Two neighboring subunits are shown as cyan Richardson diagrams.

| phi28 | Lys66 | Lys92 | Lys107 | Arg110 | Arg128 |
|---|---|---|---|---|---|
| **phi29** | n/a | n/a | Lys81 | Arg83 | Lys56 |
| **Sf6** | Arg63 | n/a | Gln81 | Arg82 | Asn101/Asn102 |
| **T4** | Lys204 | Lys83/Arg84 | Lys94/Arg95 | Lys223 | n/a |
| **P74-26** | n/a | n/a | n/a | Arg101 | Arg132 |
| **D6E** | Lys87/Arg88 | Arg96 | Lys98 | Arg101 | Lys123 |

**Table S2: Conservation of positively charged residues located in the pore.** The top row highlights positively charged residues found in the pore of the asccφ28 packaging ATPase, situated such that they can interact with substrate DNA. Superimposing solved ATPases onto the asccφ28 structure, residues proximal in cartesian space (not necessarily sequence space) are identified. If no residue is proximal, the cell is shaded red, if a polar residue is proximal the cell is shaded orange, and if a positively charged residue is proximal, the cell is shaded yellow. In some cases, more than one residue is proximal, and the cell has two residues labeled.

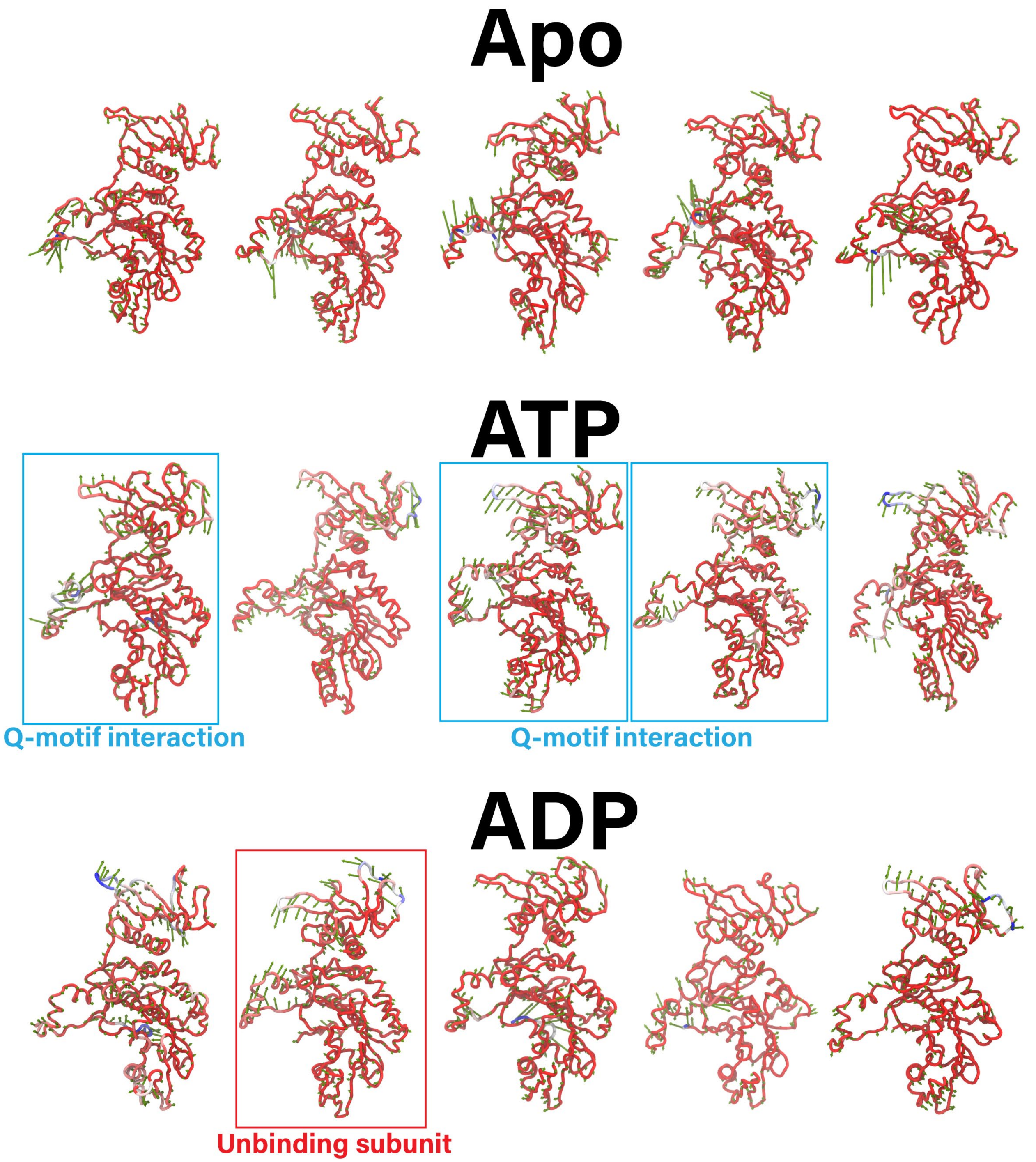


**Figure S5: the first principal component of alpha-carbon motion for all five subunits in the apo, ATP-bound, and ADP-bound states.** We find that the lid subdomain of the apo state has seemingly no correlation, suggesting that stochastic fluctuations dominate. In the ATP-bound state, the lid subdomain rotates towards the ATPase active site, particularly evident in subunits which maintain the Q-motif interaction with the adenosine base. In the ADP-bound state, the lid subdomain of the subunit which unbinds ADP rotates away from the ATPase active site. This motion is also highlighted in Movie S2. These principal components are evidence that nucleotide occupancy and binding actuate lid subdomain rotation.

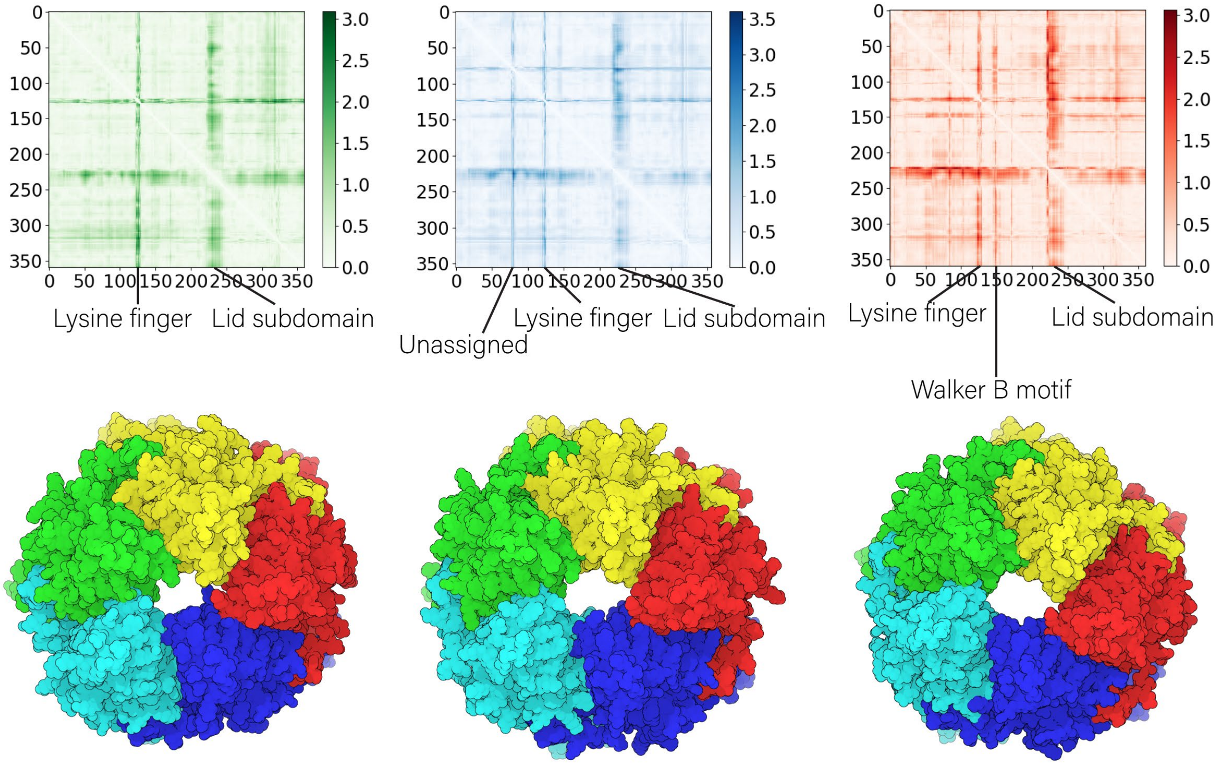


**Figure S6: variability in crystal structures.** (**Top row**) The three solved crystal structure show large standard deviations in positions of the lid subdomain and *trans*-acting lysine finger. This indicates that the lid subdomain rotation can be used to position the lysine finger of a neighboring subunit. (**Bottom row**) the crystals are not regular pentagonal structures, rather, they are oblong. This indicates that deviations in the lid subdomain, which mediated most inter-subunit contacts, can be used to change the structure's quaternary structure.

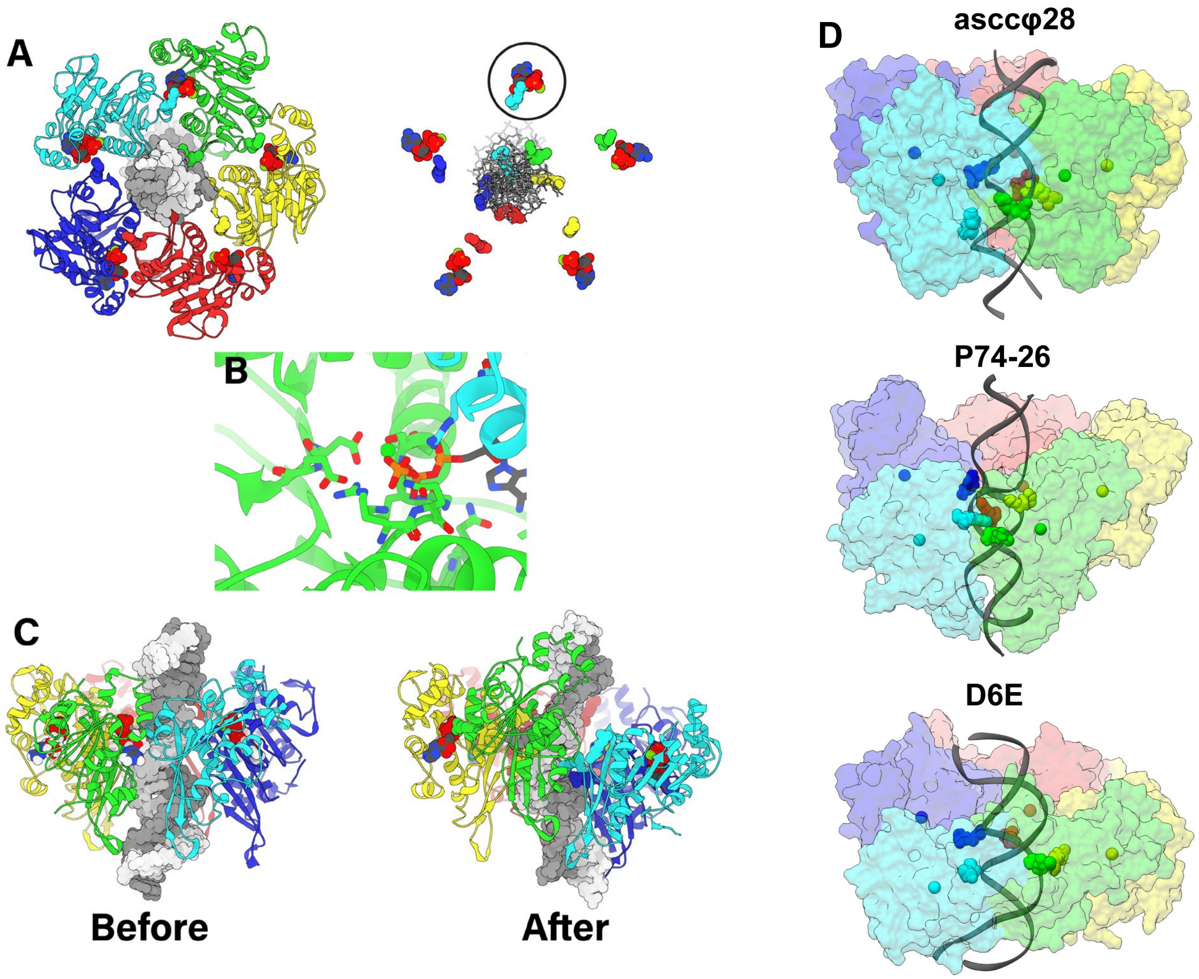


**Figure S7: P74-26 pentamer model and simulations.** (**A**) The pentamer in the presence of ATP and substrate DNA. Arg101 binds to DNA in the pore, and Arg139 interacts in trans to coordinate the γ-phosphate of ATP. These agree with biochemical data showing Arg101 is necessary to grip DNA, and Arg139 catalyzes hydrolysis in trans. On the right these interactions are shown in isolation for easier visualization. The binding pocket where Arg139 is best positioned to catalyze hydrolysis is circled. (**B**) Zooming in on a single binding pocket, where Arg139 is best positioned to catalyze ATP hydrolysis, we find that all the expected interactions are maintained based on prior monomer simulations and ATPase enzymology. (**C**) The ATP at this active site was removed and the structure was simulated in a 4-ATP-bound, 1-apo state. At the apo interface, the pentamer shears to form a helical pitch, highlighted in this before and after view. (**D**) The predicted helical arrangements of the asccφ28, P74-26, and D6E packaging ATPases are depicted. Each subunit is shown as a transparent surface. The centers of mass of each subunit are depicted as hard spheres to show the right-handed helical arrangement descending from the blue to cyan subunits. DNA-gripping residues are shown as spheres, to show their coordination with the substrate DNA, whose phosphate backbone is a dim gray ribbon. The DNA-gripping residues of asccφ28 are the same as depicted in main text **Fig. 4B**.

**Movie S1: ADP-release by the exchange residue (ER).** ADP-release is characterized by dissociation from the *cis*-acting Walker A lysine and association to the *trans*-acting EF arginine. The *cis*-acting enzyme is shown in light blue, and the *trans*-acting enzyme is shown in green. Bound $Mg^{2+}$-ADP, *cis*-lysine, *trans*-arginine, as well as residues which interact with adenosine are shown in spheres. The enzyme keeps contact with the adenosine, using the bound base as a fulcrum to pull out the phosphates.

**Movie S2: P-loop backbone conformation change upon ADP release.** As ADP is released from the subunit, the P-loop backbone dihedral angles rotate to position a backbone oxygen atom close to where the β-phosphate was prior to release.

**Movie S3: The Walker A Val30 becomes helical in absence of bound nucleotide.** In the mixed-occupancy 4-ADP-bound, 1-apo simulation, the apo enzyme Walker A motif adopts a more helical conformation. This places Val30 in the "blocking" pose observed in the iodine-derivatized crystal. This suggests that the Walker A conformation observed in this crystal is not merely an artifact of iodine-binding, but may serve a regulatory role in ADP-release and ATP-binding.

**Movie S4: Dynamics of apo monomer.** A 100-ns apo monomer simulation with the AMBER99SB-ILDN/TIP3P protein/water force field pairing. The enzyme is shown with a surface representation, and the lid subdomain is highlighted in yellow. The lid subdomain remains far from the active site and dynamic in the monomer apo simulations. This monomer result helps rationalize the observed flexibility of the lid subdomain in the apo pentamer simulation (see main text **Fig. 5**).

**Movie S5: Dynamics of ATP-bound monomer.** A 100-ns ATP-bound monomer simulation with the AMBER99SB-ILDN/TIP3P protein/water force field pairing. The enzyme is shown with a surface representation, and the lid subdomain is highlighted in yellow. Bound $Mg^{2+}$-ATP are shown as spheres. The lid subdomain rotates and closes over the binding pocket, as is expected of a lid subdomain in the ASCE superfamily. This is consistent with prior MD simulations and solved crystal structures. This monomer result helps rationalized the observed rigidity of the lid subdomain in the ATP-bound pentamer simulation (see main text **Fig. 5**). This lid subdomain rotation is proposed to be the force-generating mechanism that translocates DNA (see main text **Fig. 7**).

**Movie S6: Dynamics of ADP-bound monomer.** A 100-ns ADP-bound monomer simulation with the AMBER99SB-ILDN/TIP3P protein/water force field pairing. The enzyme is shown with a surface representation, and the lid subdomain is highlighted in yellow. Bound $Mg^{2+}$-ADP are shown as spheres. The lid subdomain rotates and closes over the binding pocket, as is expected of a lid subdomain in the ASCE superfamily. This is consistent with prior MD simulations and solved crystal structures. This monomer result helps rationalized the observed rigidity of the lid subdomain in the ADP-bound pentamer simulation (see main text **Fig. 5**). A rigid lid subdomain helps subunits which have already hydrolyzed act as a fulcrum for subunits which have not; this helps subunits not be pulled away from the viral capsid by neighboring lid subdomain rotation (see main text **Fig. 7**).

**Movie S7: D6E monomer simulation positions the lid subdomain to mediate inter-subunit contacts.** During a 2.4 microsecond simulation of the D6E monomer ATPase, the lid subdomain (yellow) extends away from the ATPase domain (green). The movie shows an interpolation between the crystal structure and the predicted extension. A gray subunit is placed by aligning both subunits to the gp11 pentamer. The interpolation shows that extending the lid subdomain would help facilitate inter-subunit contacts, akin to the gp11 pentamer.

**Movie S8: asccφ28 gp11 ATPase domain pentamer adopts a helical structure and fits into the φ29 cryo-EM reconstruction.** MD simulation of the asccφ28 gp11 pentamer ATPase domains in the 5-ATP-bound configuration predicts that the pentamer adopts a helical structure as the subunits track DNA. This pentamer fits well into the helical φ29 asymmetric cryo-EM reconstruction.

**Movie S9: P74-26 ATPase domain pentamer adopts a helical structure and fits into the φ29 cryo-EM reconstruction.** MD simulation of the P74-26 pentamer ATPase domains in the 4-ATP-bound, 1-apo configuration predicts that the pentamer adopts a helical structure as the subunits track DNA, sheared at the apo-interface. This pentamer fits well into the helical φ29 asymmetric cryo-EM reconstruction.

**Movie S10: D6E ATPase domain pentamer adopts a helical structure and fits into the φ29 cryo-EM reconstruction.** MD simulation of the D6E pentamer ATPase domains in the 5-ATP-bound configuration predicts that the pentamer adopts a helical structure as the subunits track DNA. This pentamer fits well into the helical φ29 asymmetric cryo-EM reconstruction.
