## Supplemental Movies for "Atomistic Basis of Force Generation, Translocation, and Coordination in a Viral Genome Packaging Motor"

### Slide 1
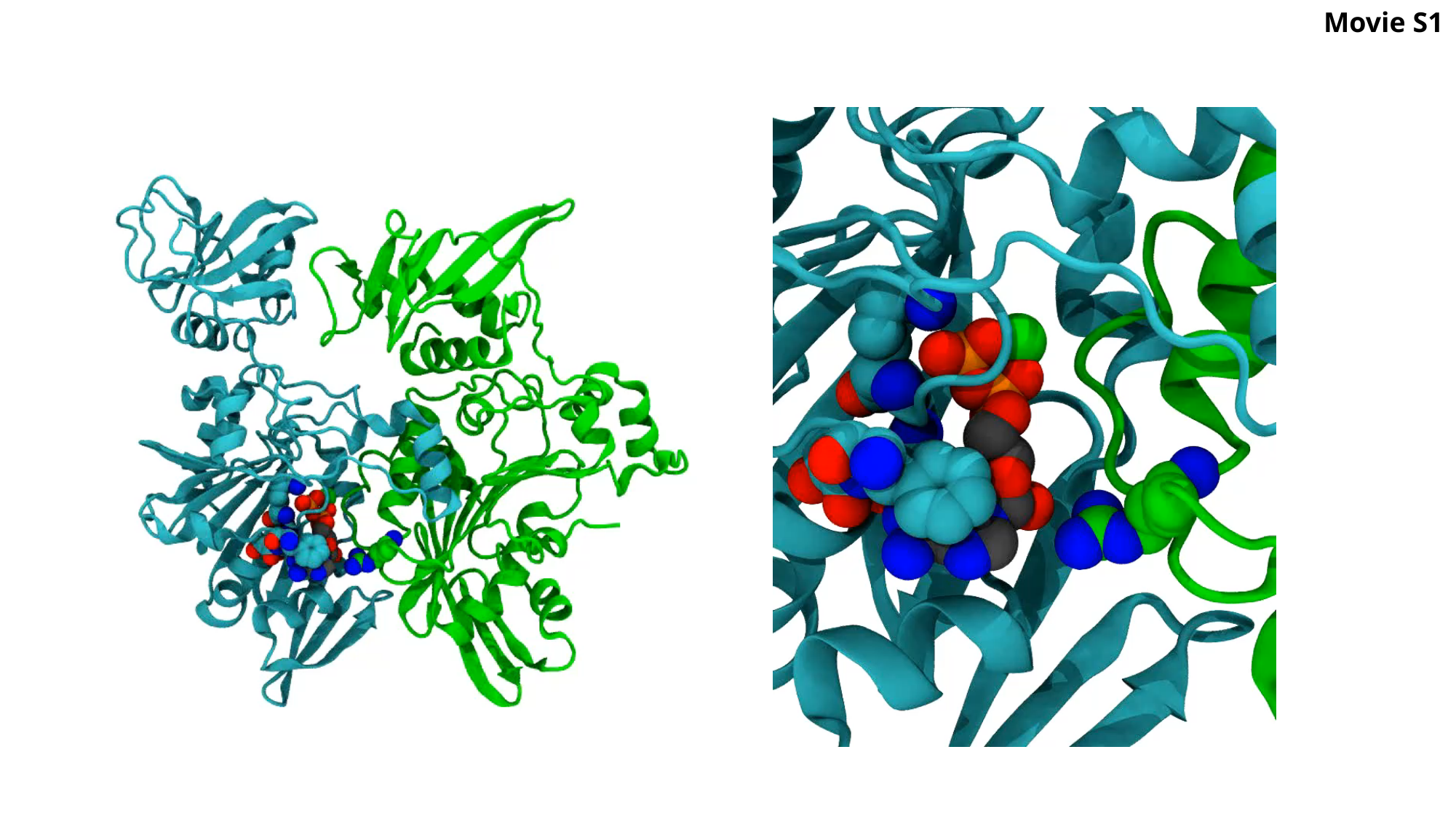

Movie S1

### Slide 2
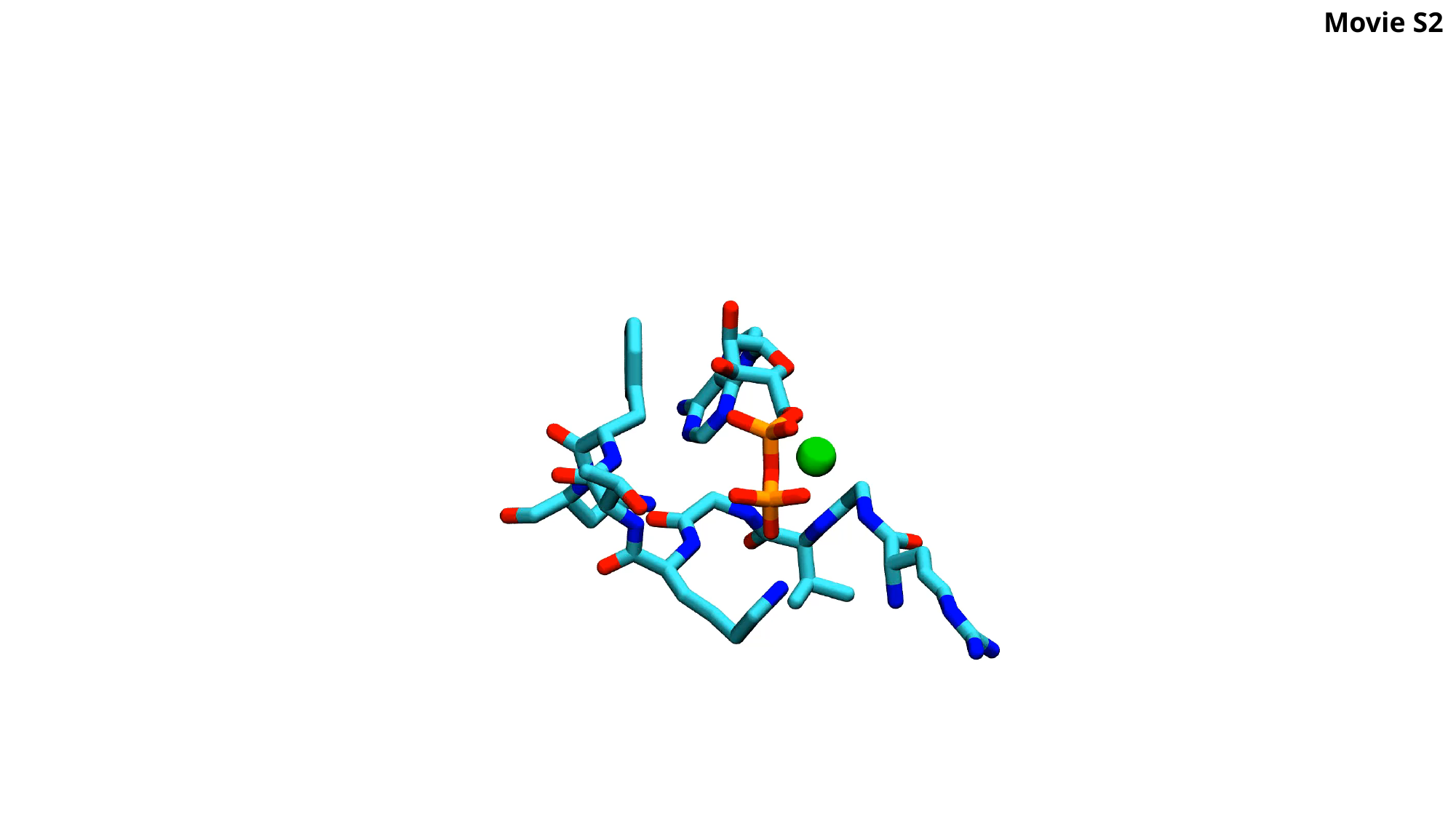

Movie S2
#

### Slide 3
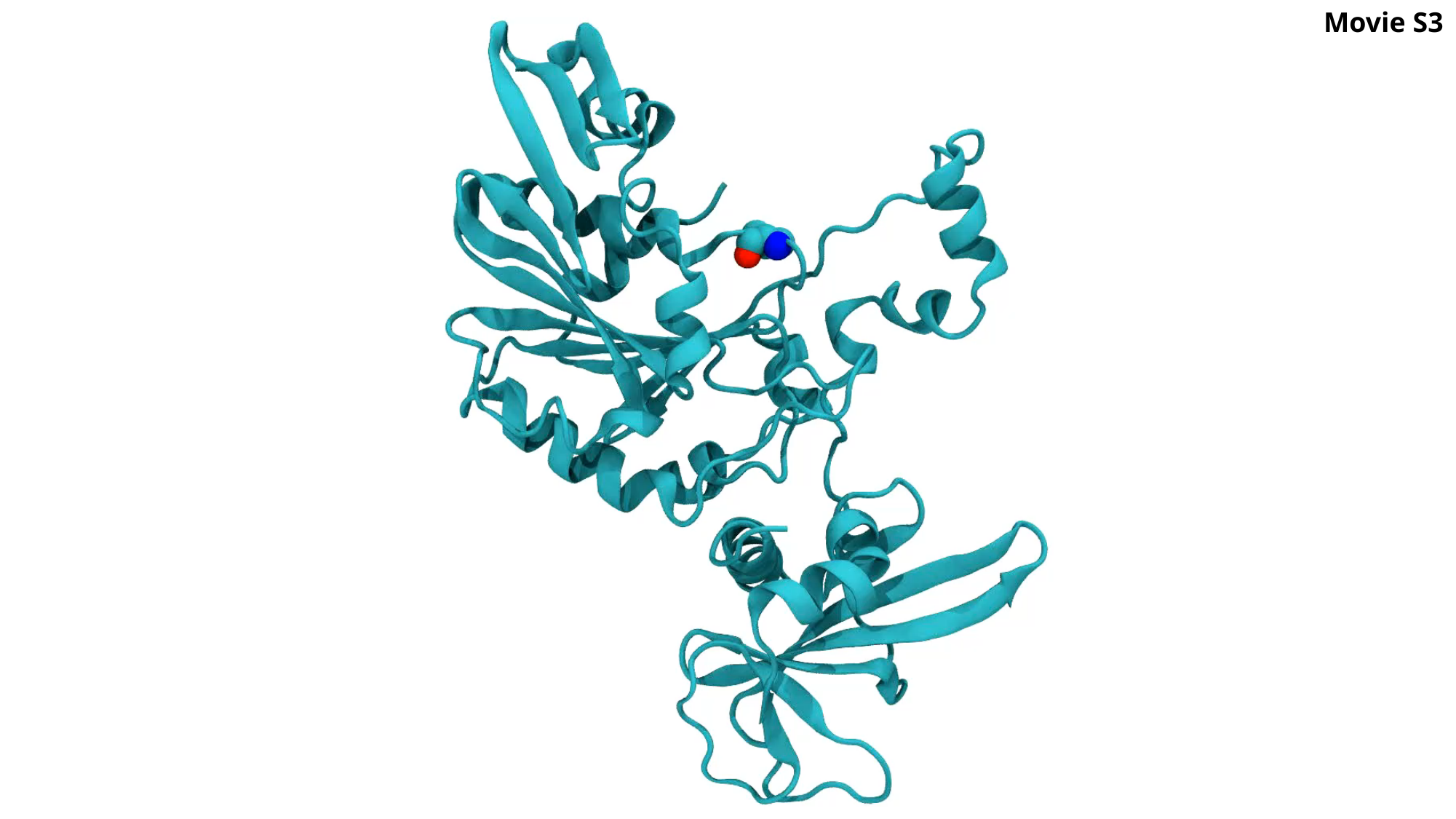

Movie S3
#

### Slide 4
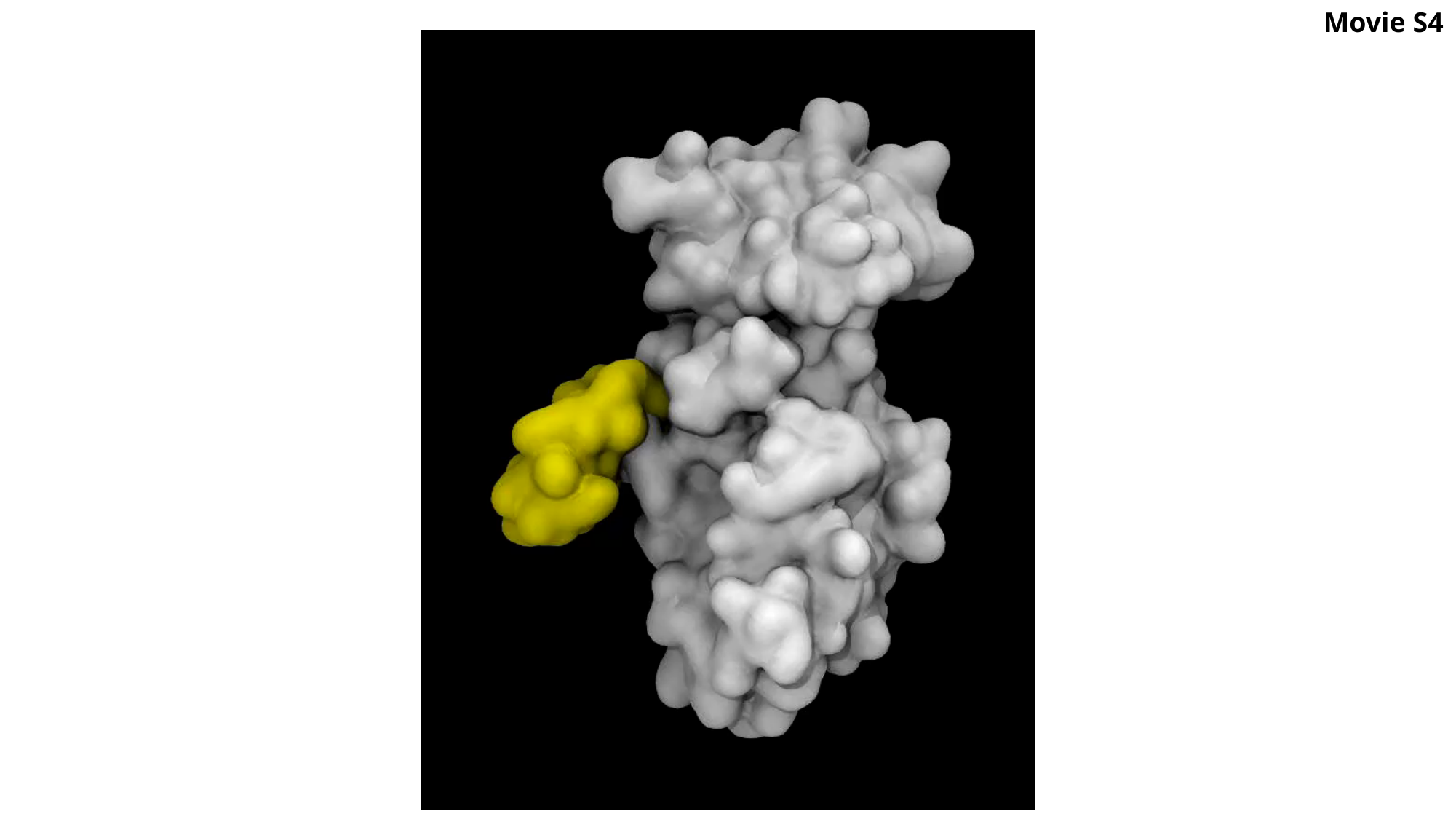

Movie S4
#

### Slide 5
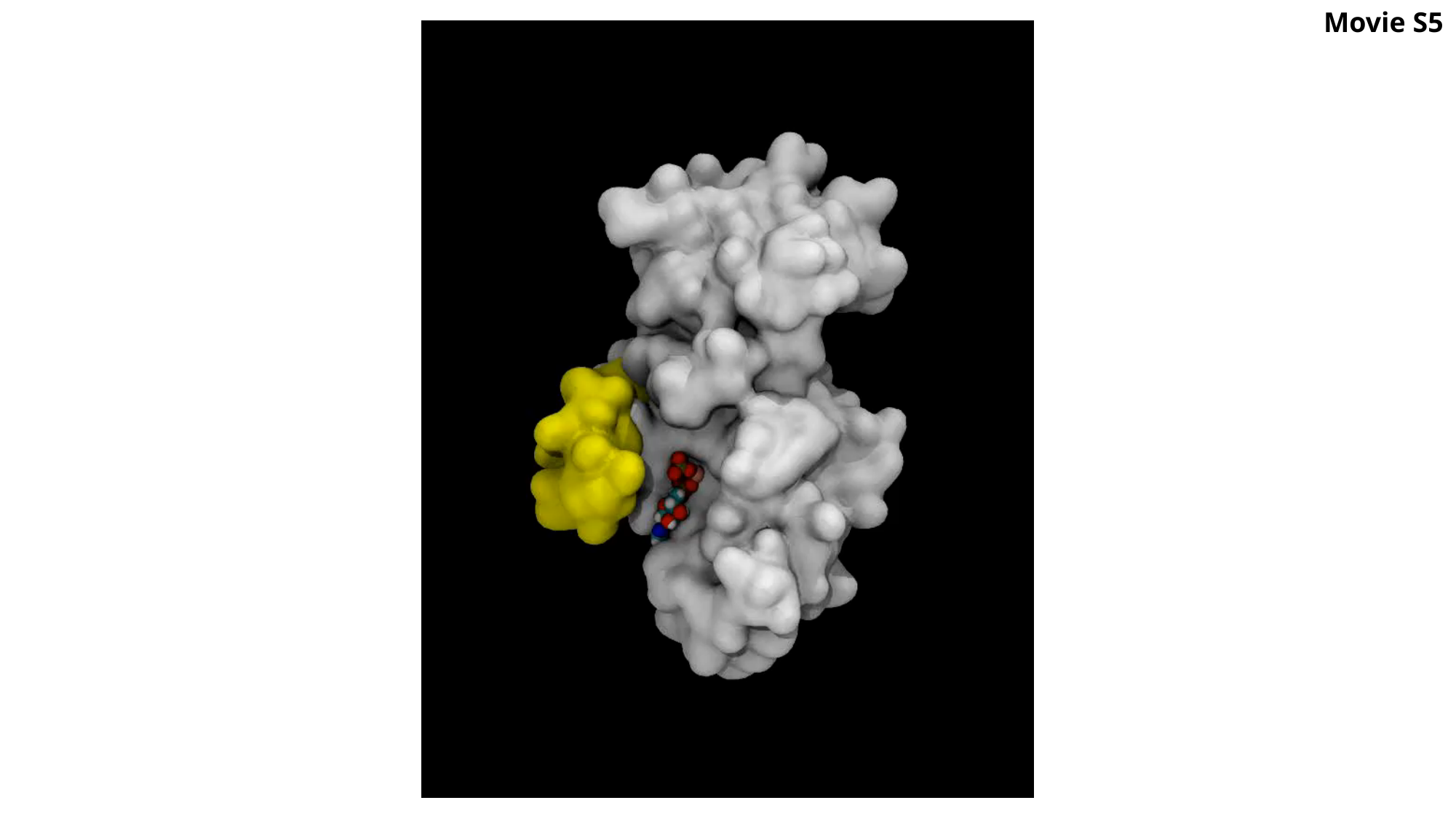

Movie S5
#

### Slide 6
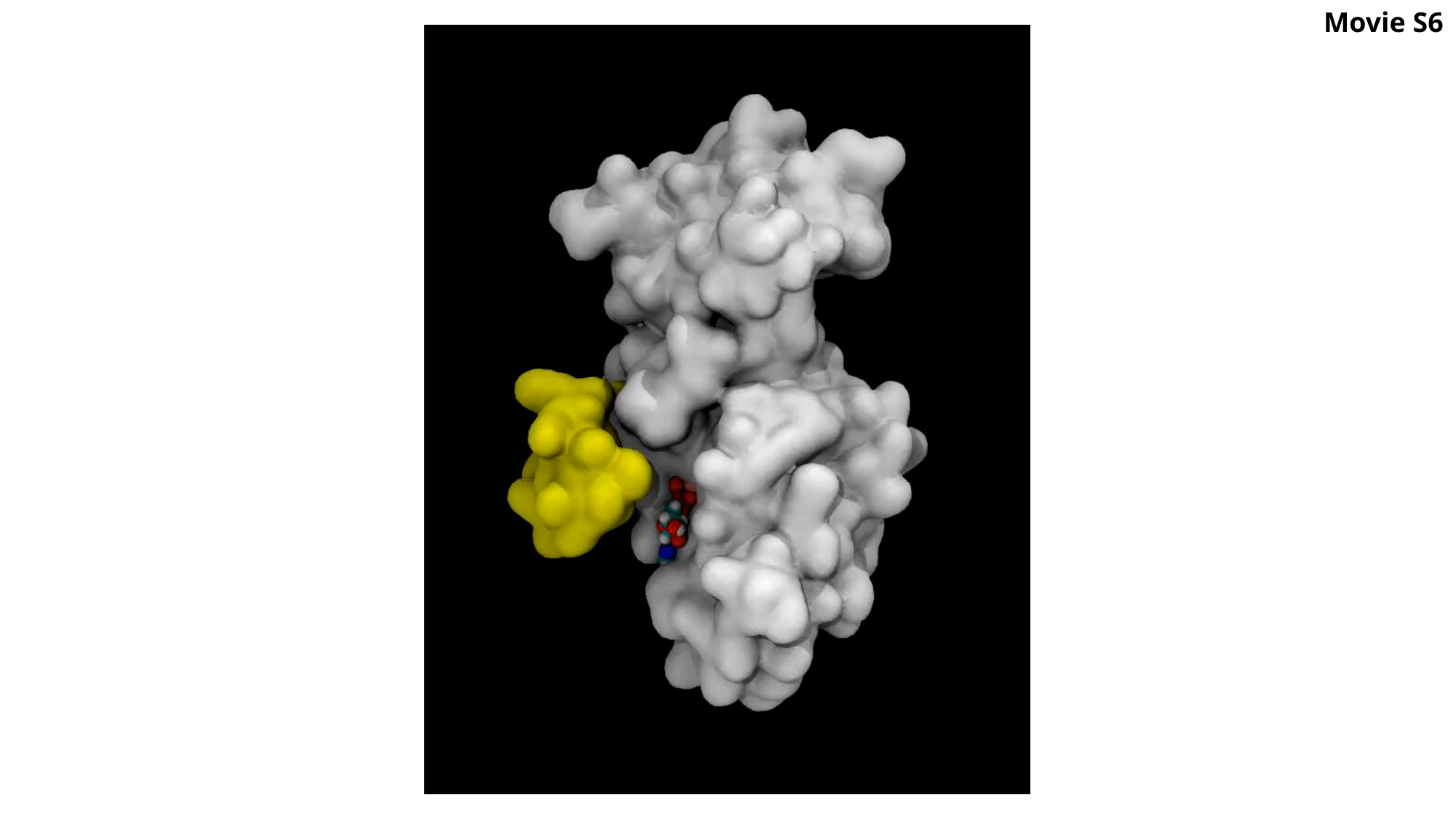

Movie S6
#

### Slide 7
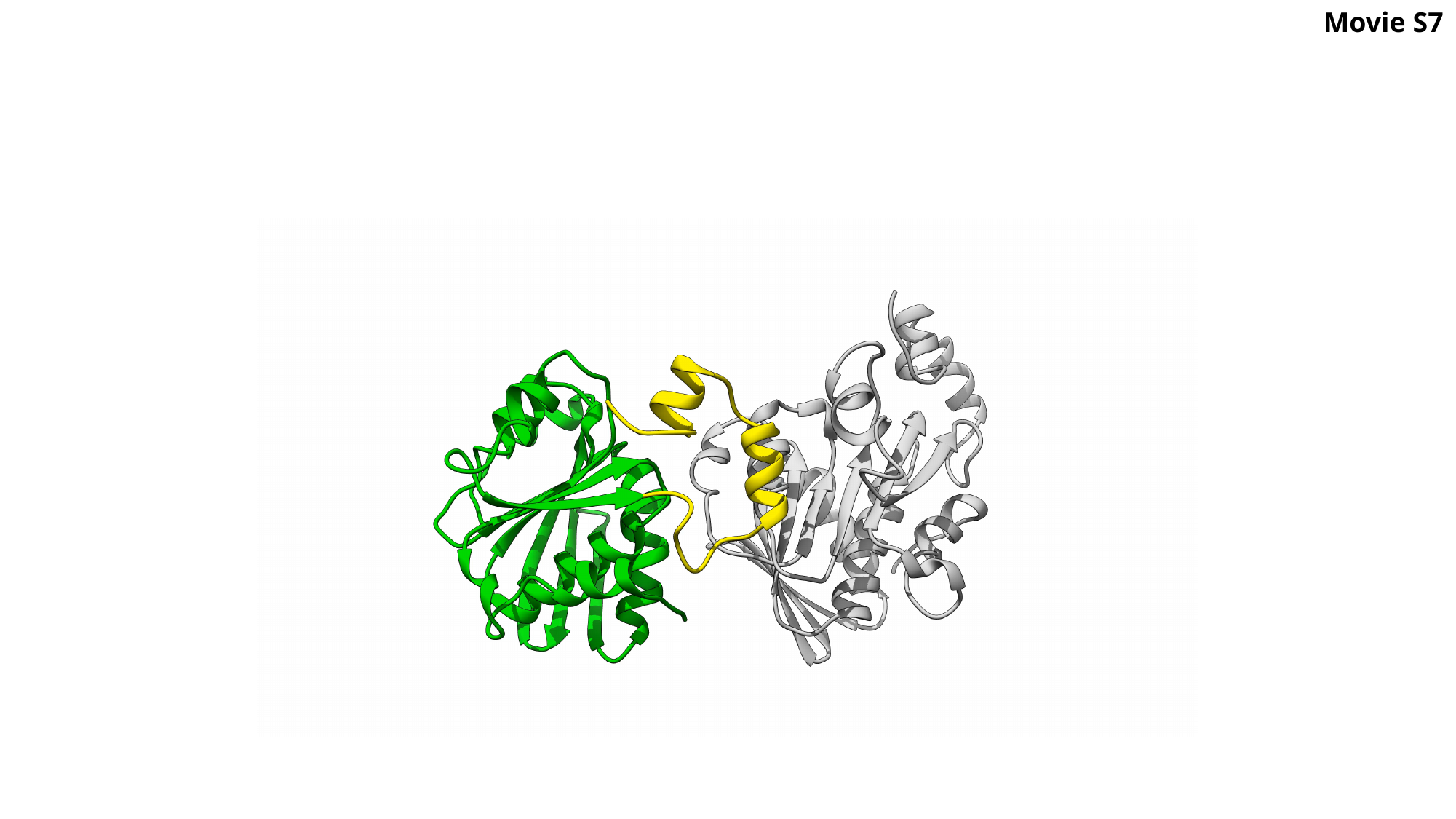

Movie S7
#

### Slide 8
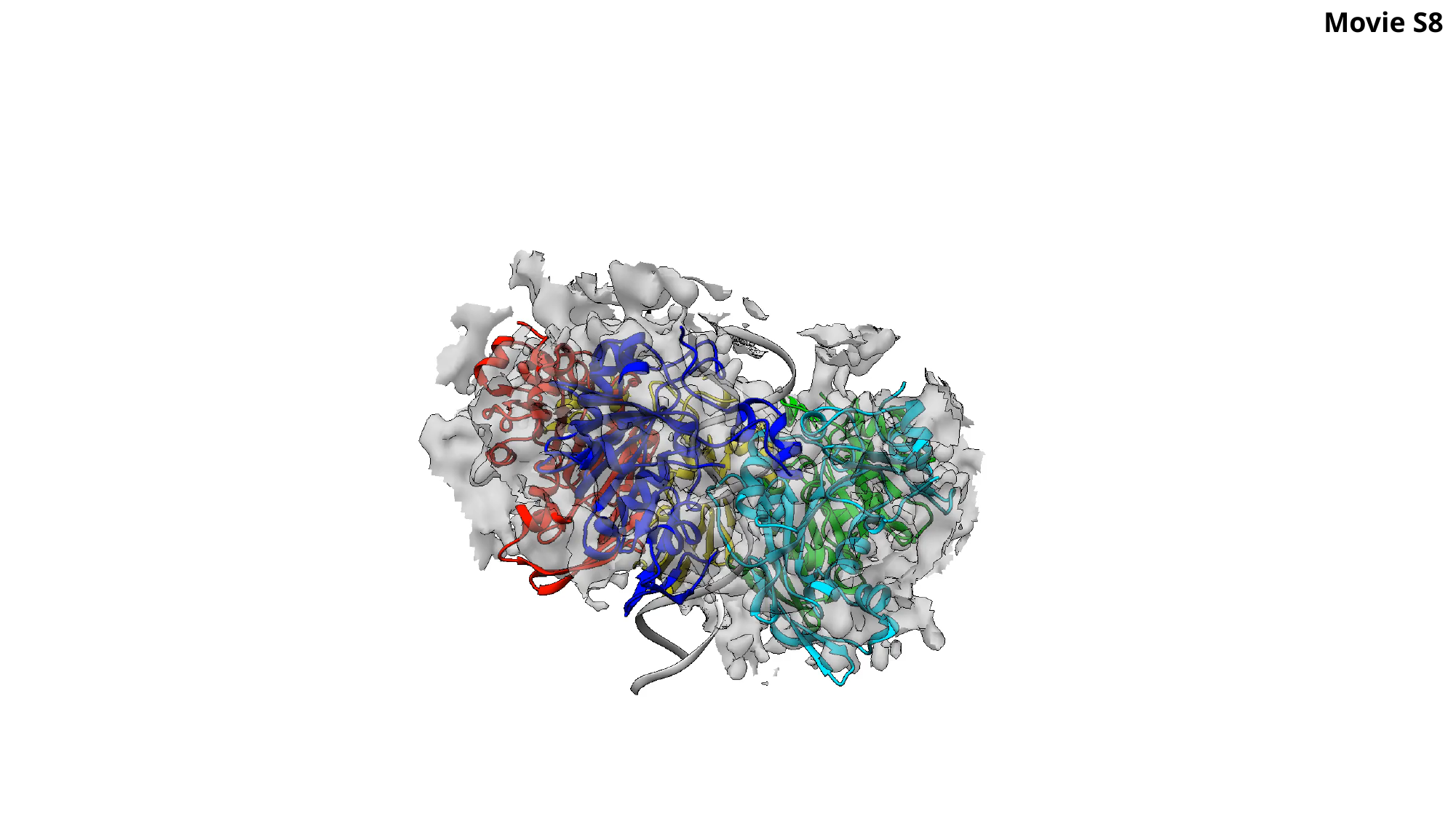

Movie S8
#

### Slide 9
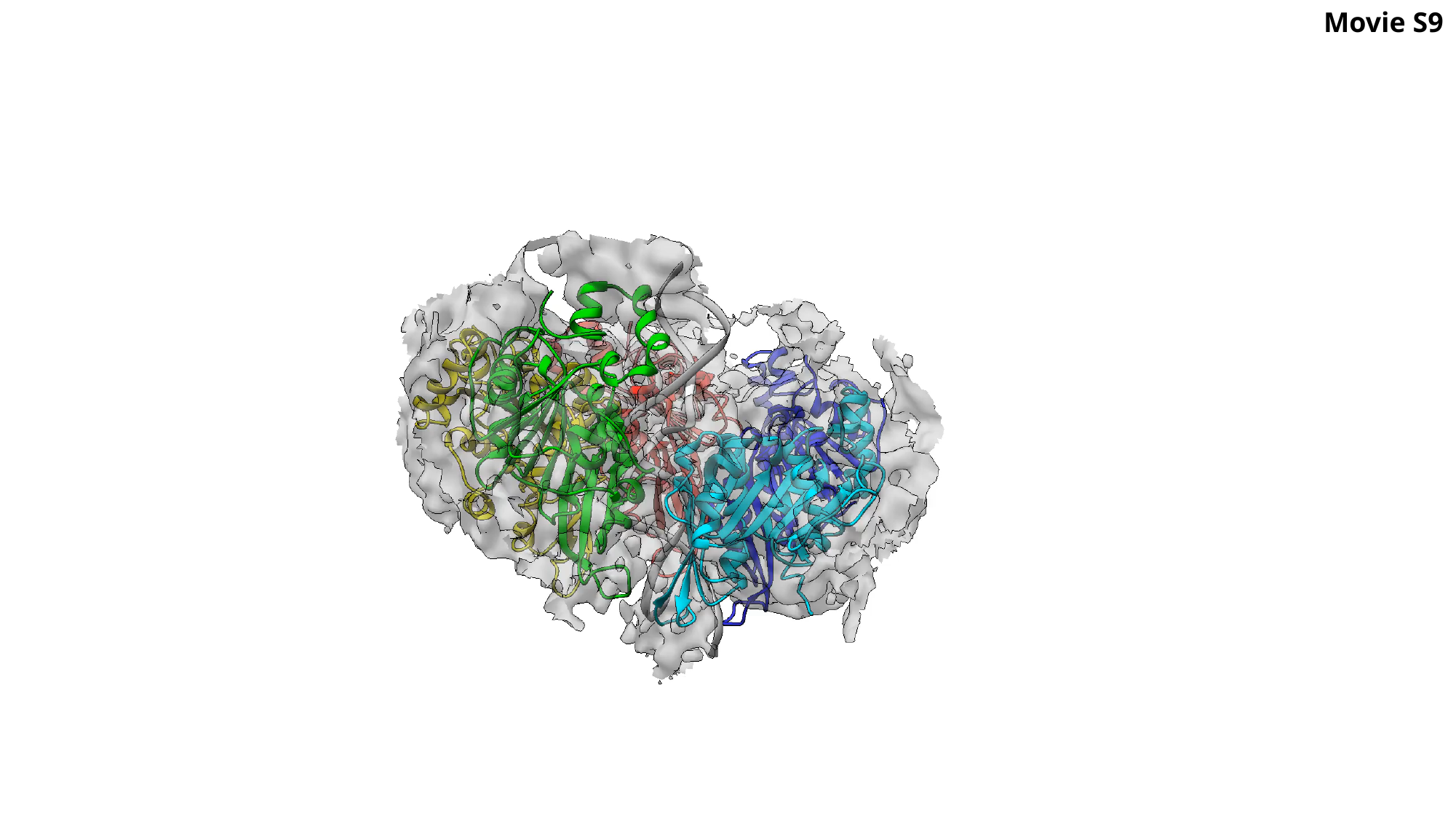

Movie S9
#

### Slide 10
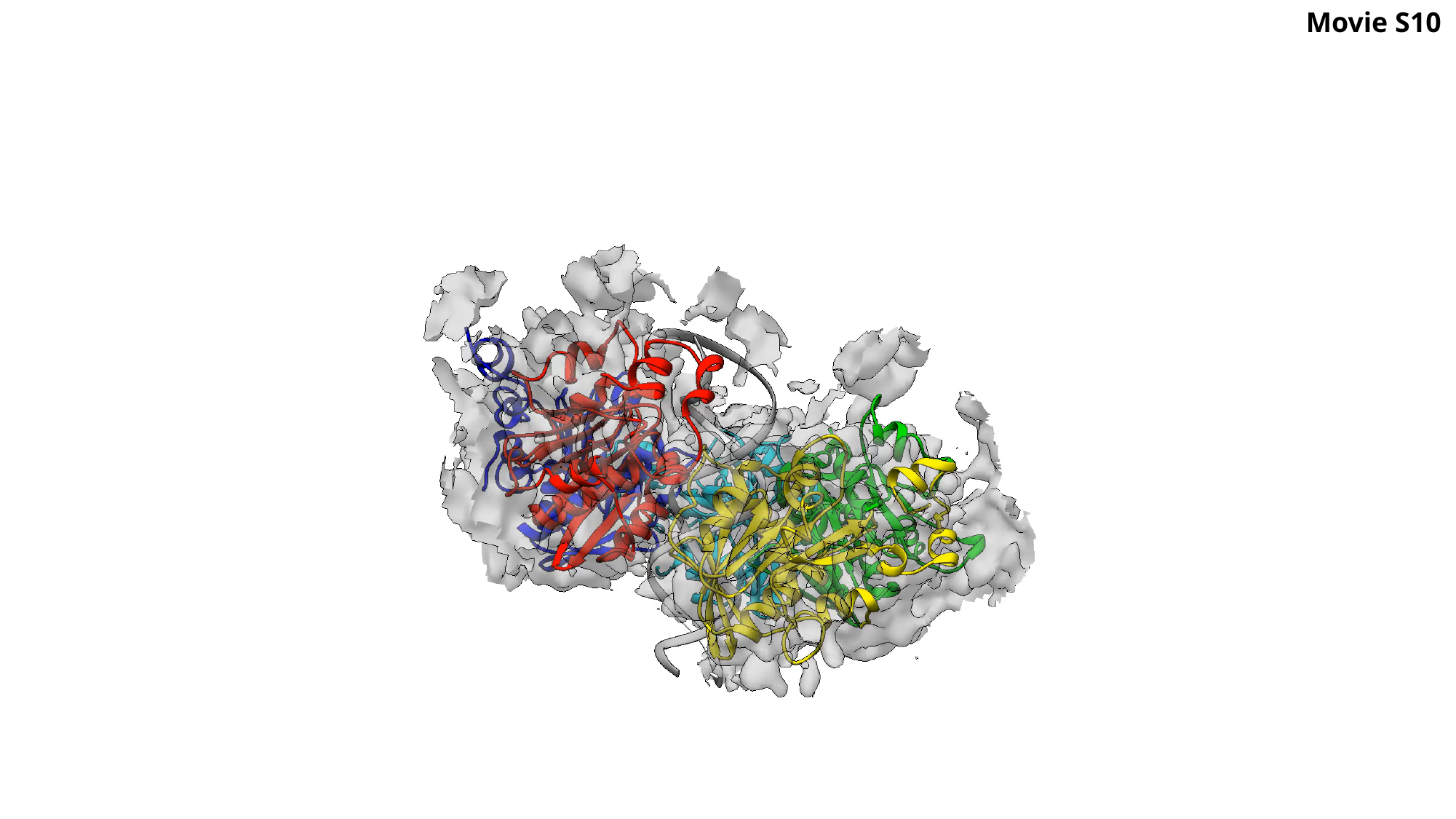

Movie S10
#
